# Long-term responses of life-history strategies to climatic variability in flowering plants

**DOI:** 10.1101/2022.10.19.512857

**Authors:** James D. Boyko, Eric R. Hagen, Jeremy M. Beaulieu, Thais Vasconcelos

## Abstract

- The evolution of annual or perennial strategies in flowering plants may depend on a broad array of temperature and precipitation variables. Previously documented correlations between life history strategy and climate appear to be clade-specific and fail to consider the coevolution of climatic niches and life history strategies.
- Here we combine annual and perennial life history data with geographic distribution for 9,939 flowering plant species and utilize a recently developed method that accounts for the joint evolution of continuous and discrete traits to evaluate two hypotheses: (1) annuals tend to evolve in highly seasonal regions prone to extreme heat and drought, and (2) annuals tend to have faster rates of climatic niche evolution than perennials.
- We find temperature, particularly the maximum temperature of the warmest month, is the most consistent climatic factor influencing life history evolution in flowering plants. Unexpectedly, we find that the rates of climatic niche evolution are faster in perennials than in annual lineages.
- We propose that annuals are consistently favored in areas prone to extreme heat due to their ability to escape heat stress as seeds, but they tend to be outcompeted by perennials in regions where extreme heat is uncommon or nonexistent.

## Introduction

Flowering plants have evolved multiple different life forms and life history strategies to survive environmental challenges (Grime 1977; Stearns 1992). For instance, resprouting plants can regenerate after fire or drought from dormant underground buds (e.g. Rando et al. 2016; Howard et al. 2019), and large trees can annually shed their leaves or protect their growing buds during freezing conditions with scale-like modified leaves (Raunkiaer 1934; Zanne et al. 2014; Edwards et al. 2017). Other plants have increasingly shortened their life cycles so that germination, fertilization, and seed release all happen within the favorable season of a single year, allowing them to avoid the unfavorable season in the form of seeds (Mulroy and Rundel 1977; Zanne et al. 2014). The latter describes the life history strategy of annual plants, which are semelparous (i.e., reproduce just once before death; Stearns 1992). This is opposite to the vast majority of flowering plant species, which are mostly iteroparous (i.e., present multiple reproductive events) and characterized by a perennial life history strategy with adaptations to survive an indefinite number of unfavorable seasons (Raunkiaer 1934; Friedman 2020).

There is an uneven distribution of annual and perennial strategies throughout the globe, and this observation has generated interest in finding environmental correlates associated with the evolution of different life history strategies in flowering plants (Figure 1; Raunkiaer 1934; Ricklefs and Renner 1994; Friedman 2020). Perennial plants have a bimodal distribution of diversity with peaks in warmer tropical climates (Grime 1977) and areas where freezing is constant, such as higher latitudes and alpine habitats (Billings and Mooney 1968; Givnish 2015). By contrast, annuals are highly represented in mid-latitude areas subject to prolonged drought, such as desert and Mediterranean habitats (Mulroy and Rundel 1977). Annual angiosperms can represent over half of the floristic diversity in these regions (Figure 1b; Raunkiaer 1934) despite being considerably less common than perennials on a global scale (Friedman 2020).

**Figure 1.**
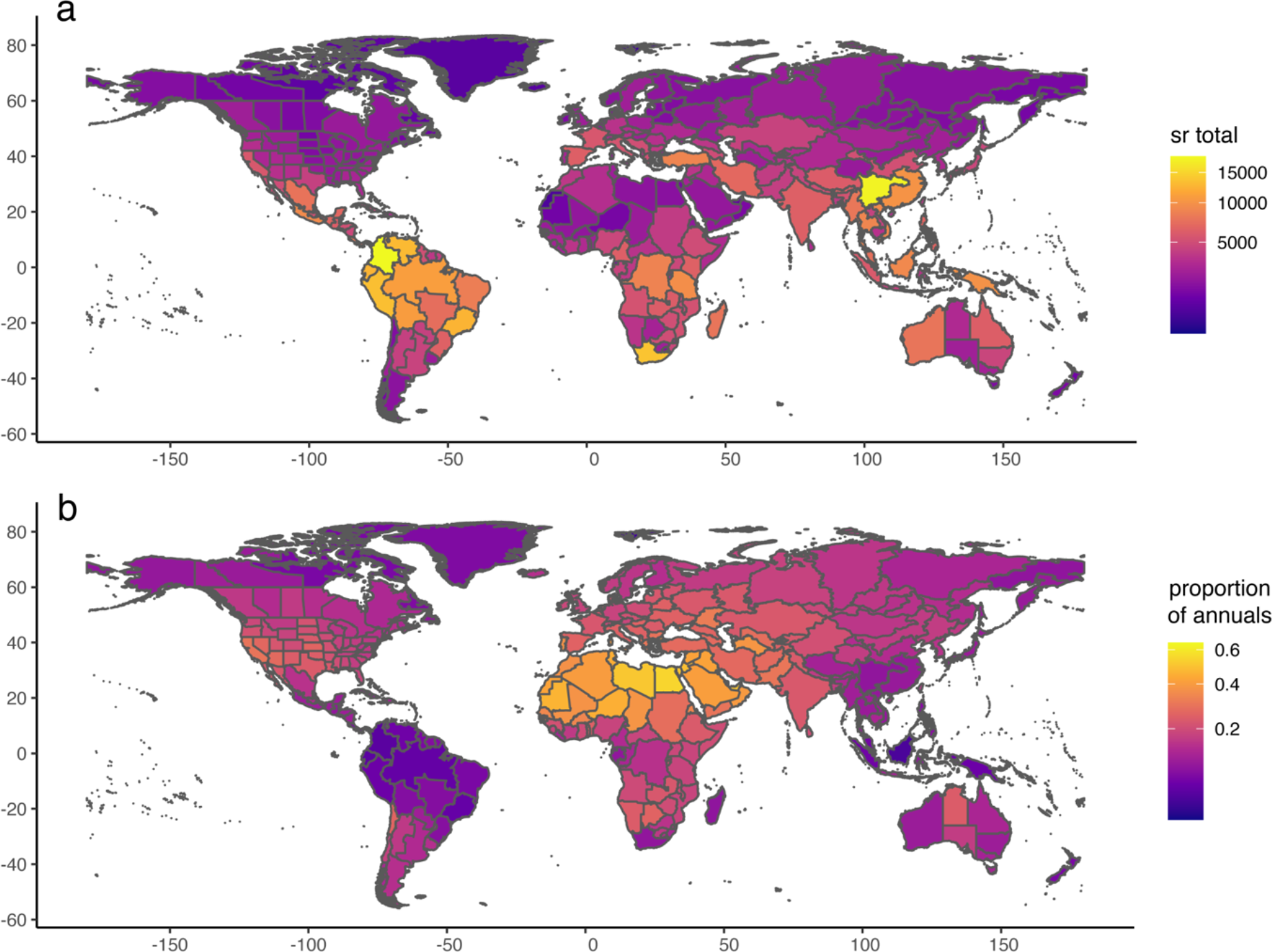
Global distribution of vascular plant diversity and proportion of annual plants. (a) Total species richness of vascular plants by botanical country according to the WCVP database (WCVP, 2022), and (b) Proportion of annual plants in relation to total species richness. Y-axis: longitude; x-axis: latitude.

Although the uneven distribution of different life forms across the globe has long been recognized (Raunkiaer 1934; Stebbins 1974; Grime 1977; Friedman 2020), the historical drivers of this pattern are still debated, and much of the discussion has focused on the role of climate. For instance, according to the theory of life history strategies in plants, annuals are more likely to evolve where the climate is seasonal because they can rapidly take advantage of short beneficial climatic conditions for reproduction (Cole 1954; Friedman 2020). Support for this has been found in clades typical of Mediterranean habitats, such as *Heliophila* (Brassicaceae) in Africa (Monroe et al. 2019), *Bellis* (Asteraceae) in Europe (Fiz et al. 2002), and grasses (Poaceae) across the globe (Humphreys and Linder 2013). Others have argued that the evolution of the annual life form is linked to the occupation of generally warmer environments (Stearns 1992), and support for this has been found in temperate clades such as Montiaceae (Ogburn and Edwards 2015). Similarly, annuals might frequently be excluded from alpine environments where frost is common due to high seedling mortality (Givnish 2015). Finally, some have argued that temperature and precipitation, as well as their annual seasonality, are relevant in explaining the evolution of different strategies, as shown in *Oenothera* (Onagraceae; Evans et al. 2005). Despite disagreements about the specific climatic controls on the biogeography of life history strategies, there is a clear consensus that both temperature and precipitation likely influence these patterns. However, empirical studies aiming to correlate climate with life history distributions have so far focused on specific clades or geographic areas. It is unclear which relationships are sufficiently general to be consistent when multiple clades are considered in the same analytical framework.

In addition to general climatic preferences, previous work has not thoroughly explored how different life history strategies may affect the ability to adapt to changing climatic circumstances. Once a life history strategy has evolved in response to particular climatic pressures, it may also impact long-term biogeographical patterns of lineages that possess it. For example, the evolution of the annual habit is linked to a series of traits associated with more secure reproduction and increased vagility, such as selfing (Stebbins 1950; Aarssen 2000) and relatively high investment in seed production (Friedman 2020). For these reasons, annuals are considered to be generally good invaders (Pannell et al. 2015; Linder et al. 2018). Furthermore, due to their generally shorter generation times, annuals may also exhibit faster rates of phenotypic evolution (e.g., Smith and Beaulieu 2009), possibly allowing them to adapt more quickly to changing environmental conditions (Andreasen and Baldwin 2001).

Here, we assess the relationship between climatic factors and the evolution of life history strategies in flowering plants. To that end, we apply recent developments in trait evolution models (Boyko et al. 2022) to explicitly incorporate climatic niche variation’s impact on life history strategy evolution. We account for the heterogeneity of evolutionary histories in flowering plants and the habitats associated with them by analyzing a broad sample of clades with global distribution and where multiple transitions between annual and perennial strategies are observed. Two specific hypotheses are addressed: (1) annuals evolve in warmer and drier climates, or where seasonality is stronger, more frequently than perennials, and (2) annuals tend to have faster rates of climatic niche evolution than perennials due to their shorter generation times and propensity to establish themselves in new environments. We expect to find mixed support for our hypotheses due to clade-specific evolutionary patterns. Some clades will undoubtedly have more heterogeneity in transition rates between life history strategies, whereas other clades may have exclusively unidirectional transitions, and others still may have no heterogeneity at all. However, due to our large dataset and the ability to account for rate heterogeneity in our models, we expect that we can illuminate the generalities of the long-term responses of life-history strategies to climatic variability in flowering plants.

## Materials and Methods

### Phylogenetic and life history datasets

To build a dataset of life history strategies for a set of flowering plant clades, we used the recent release of the World Checklist of Vascular Plants dataset (WCVP 2022), which includes life form data following the Raunkiaer (1934) system. The Raunkiaer system classifies different life history strategies in flowering plants based on the position of the buds in relation to the soil at the end of the growing season as well as on how plants protect growing buds during the unfavorable seasons. We scored as “annuals” all species marked as “Therophytes” (including combinations such as “Climbing therophyte” and “Semiaquatic therophyte”) in the WCVP dataset. All other life forms, such as “Biennials,” “Cryptophytes,” “Nanophanerophytes,” and “Phanerophytes” were scored as “perennials.”

Following this scoring, the proportion of “annuals” to “perennials” in the WCVP dataset is around 1:4. Annual plants are considerably less common than perennials, and it is more common to find clades where all species are perennials than clades where evolutionary transitions between annual and perennial strategies are observed. However, we restricted our set of clades to groups that presented multiple evolutionary transitions between different annual and perennial life history strategies. Selecting only groups where both life history states are present will certainly bias our view of how different life histories and climatic niches impact each other over evolutionary time. Our analytical framework partially mitigates this bias by accounting for hidden heterogeneity arising from character-independent continuous trait evolution.

The set of clades selected for our analyses is not restricted to a single taxonomic rank and includes any clade that matched these criteria: (1) both annuals and perennial strategies are observed, (2) a time-calibrated phylogenetic tree is available in the literature, and (3) the phylogeny includes 50 to 1000 tips and at least 10% of the known species diversity in the clade. The selected clades and the sources of their phylogenies are depicted in Table 1. In total, our study includes 32 phylogenetic trees with a total of 9,939 taxa that are distributed globally (Table 1). We also completed the life form scoring by adding data collected from the literature so that each clade had a maximum of 30% missing data.

**Table 1.**
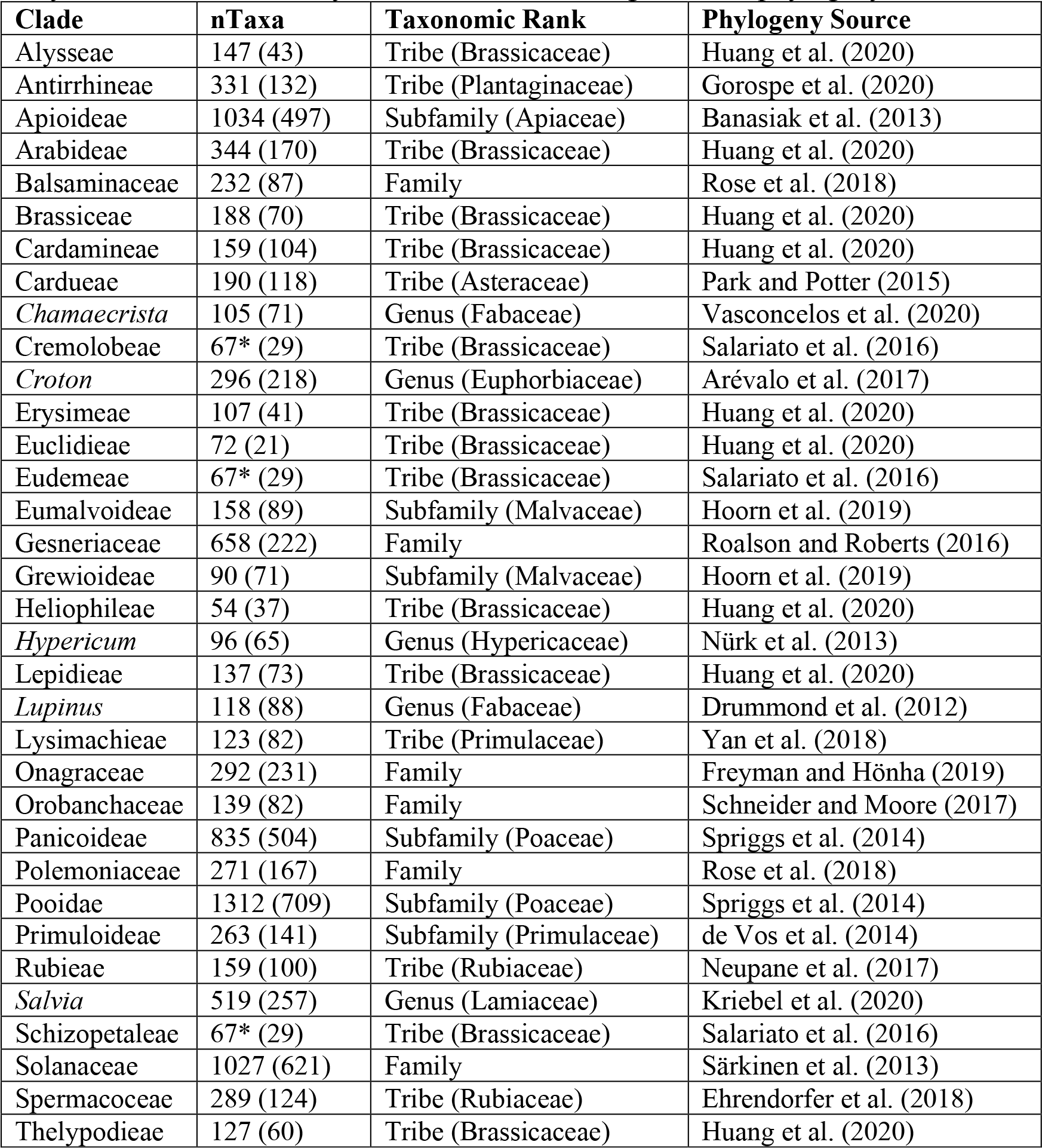
The 32 clades used in our analysis as well as their taxonomic rank, source, and size of the phylogeny. Numbers in brackets are the number of taxa with available data used in analysis. Clades indicated by * are members of a single “CES” phylogeny.

### Distribution points and climatic data

We standardized all species names in the phylogenies following the GBIF taxonomic backbone with the R package *taxize* (Chamberlain and Szöcs 2013) and then downloaded occurrence points that had preserved specimens associated with these names using functions of the R package *rgbif* (Chamberlain and Boettiger 2017). This resulted in a dataset of 3,155,956 occurrence points. We filtered the points according to the native distribution range of genera and species using the shapefiles of the Working Group on Taxonomic Databases for Plant Sciences (TDWG) for level 3 botanical countries (Brummitt et al. 2001) combined with the WCVP dataset. Filtering was particularly important to exclude the invasive ranges of several species, keeping only native ranges according to the expertise of taxonomists (POWO 2022). Other irregularities such as points in the sea, outliers, duplicated coordinates for the same species, and centroids of countries were also removed using a protocol similar to the one described in Vasconcelos et al. (2021).

We examined seven climatic variables from CHELSA (Climatologies at high resolution for the earth’s land surface areas; Karger et al. 2017; Table 2): BIO 1: Mean Annual Temperature (MAT), BIO 4: Temperature Seasonality, BIO5: Maximum Temperature of the Warmest Month, BIO6: Minimum Temperature of the Coldest Month, BIO 12: Mean Annual Precipitation (MAP), BIO15: Precipitation Seasonality, and BIO17: Precipitation of Driest Quarter. Aridity Index (AI), where higher values indicate greater humidity, was also included in the analysis (Trabucco and Zomer 2019). All variables were analyzed at the resolution of 30 arc-seconds (1 km at the equator). To summarize climatic data for each species, we used functions of the R packages *sp* and *raster* (Bivand et al. 2008; Hijmans et al. 2015) to extract a value for each filtered occurrence point based on the climatic layers we assembled. To mitigate the impact of collecting bias, we filtered these points so that no more than one occurrence point for every 1 by 1 degree cell for each species was included. The value of each remaining point was then log-transformed and used to calculate mean and within-species variance (Labra et al. 2009) for each species, the latter of which was used as error measurement in downstream analyses.

**Table 2.** Inequalities describing how expected values and expected variances will differ for each climatic variable. When Annual > Perennial, we hypothesize that the climatic niche value for that variable will be greater for annuals than perennials. When Perennial > Annual, we expect the climatic niche value for that variable to be greater for perennials than annuals. For all variables, we expect annuals to present higher rates of climatic niche evolution (i.e., higher expected variance) for annuals than perennials.

| Climatic Variable | Expected Value | Expected Variance |
| --- | --- | --- |
| Annual Mean Temperature | Annual > Perennial | Annual > Perennial |
| Temperature Seasonality | Annual > Perennial | Annual > Perennial |
| Max Temperature of Warmest Month | Annual > Perennial | Annual > Perennial |
| Min Temperature of Coldest Month | Annual > Perennial | Annual > Perennial |
| Annual Precipitation | Perennial > Annual | Annual > Perennial |
| Precipitation of Driest Month | Perennial > Annual | Annual > Perennial |
| Precipitation Seasonality | Annual > Perennial | Annual > Perennial |
| Aridity Index | Perennial > Annual | Annual > Perennial |

### Trait evolution analyses

Our analysis was conducted with two complementary goals in mind. First, we wanted to accurately model correlations between climatic niche evolution and life history characters within each of our 32 clades. This was done by fitting a set of 15 *hOUwie* models with 100 stochastic mappings per iteration and adaptive sampling enabled (Boyko et al. 2022). *hOUwie* is a recently developed framework that explicitly models the joint evolution of discrete and continuous characters. Each of the fitted model structures can be parameterized such that the evolution of the continuous trait and the discrete character are correlated (character-dependent) or uncorrelated (character-independent). In the context of our analyses, the character-dependent models test for an explicit difference in climatic niche evolution between annual and perennial lineages, whereas character-independent model structures assume no difference. Furthermore, several models have a mixture of character-dependent and character-independent processes, allowing some of the model’s parameters to depend on life history while others are fixed as equal. For example, a model which allows the rate of climatic niche evolution to vary depending on whether a lineage is annual or perennial, while also fixing their climatic optima to be equal, would mix character-dependence and character-independence. Finally, we included completely character-independent models that allow for rate heterogeneity independent of the focal trait. These types of models are important as null hypotheses which account for the possibility that our model selection is biased towards correlation as a consequence of detecting rate heterogeneity without true correlation (Beaulieu and O’Meara 2016; Caetano et al. 2018; Boyko and Beaulieu 2022). These models account for the fact that climatic niche evolution is likely to be variable throughout the phylogeny regardless of potential correlation with life history.

In total, we fit six character-independent models (CID), four character-dependent models (CD), and four hybrid models (HYB) that include both character-dependent and character-independent rate heterogeneity. The parameters we allowed to vary in our model are rates of transition between annual and perennial (*q*), the phenotypic optima of the climatic niche (*θ*), and the rate of climatic niche evolution (*σ*^2^). This means we analyzed BMV, OUV, OUM, and OUMV type models, as well as BM1 and OU1 (Boyko et al. 2022). We conducted model-averaging and compared parameter estimates within *hOUwie* to test for: (1) a relationship between climatic optima and life history strategy, and (2) whether evolutionary rates of annuals are greater than those of perennials across all climatic variables. However, rather than comparing parameter estimates (*θ*, *σ*^2^, *q*) directly, we compared the expected values and expected variances of the tips, which combine the parameter estimates and the phylogenetic history of each lineage (Hansen 1997; Butler and King 2004; Beaulieu et al. 2012). The value of a parameter estimate in isolation can be misleading because its interpretation will depend on the value of other aspects of the model. For example, although *θ* indicates a long-term phenotypic optimum, the speed at which that optimum is approached as well as the biological significance of that estimate will depend on the amount of time spent in a particular state (*q*) and the rate of pull towards the optimum (*α*) while in that state. By using the expected value and expected variance, we can evaluate whether the model predicts differences between annuals and perennials while accounting for all the model parameters and the inherent uncertainty in the evolutionary histories of the lineages.

The differences between expected values and expected variance between annuals and perennials were hypothesized to depend on the particular climatic variable being modeled (Table 2). For each clade, we tested whether there was a signal of correlation between the climatic variable and life history strategy by comparing the AIC values for our different model types (CD, CID, and HYB). A value quantifying overall model set support for correlation was calculated as the sum of the AICc weights for the model multiplied by either 1 for CD class models, 0 for CID class models, or 0.5 for HYB models. Each model set was applied independently to the 32 clades and their 8 climate variable datasets. Following the model fitting, we model-averaged each tip’s expected value and variance by the AIC weight of the model fit with which it is associated. These tip values were then categorized as either annual or perennial, and the mean of each discrete category was taken for each clade. Each tip will always have the same observed state (unless explicitly coded as unknown), but their hidden state may differ. Thus, all estimated parameters were averaged over hidden rate classes based on the associated observed character and joint probability of the underlying regime.

The final part of our analysis tested whether the associations we detect within clades are broadly consistent across the 32 clades. We used paired t-tests that incorporated phylogenetic information (Revell 2012) to assess whether model-averaged expected values and variances associated with life history strategy are consistently different across several clades. We used the whole seed plant phylogeny based on molecular data from Smith and Brown (2018; “GBMB” tree) as a template to generate a backbone phylogeny that includes each of the 32 clades as individual tips (Figure 2a), using the R packages *phangorn* (Schliep 2011) and *ape* (Paradis et al. 2004) to prune out all other tips.

**Figure 2.**
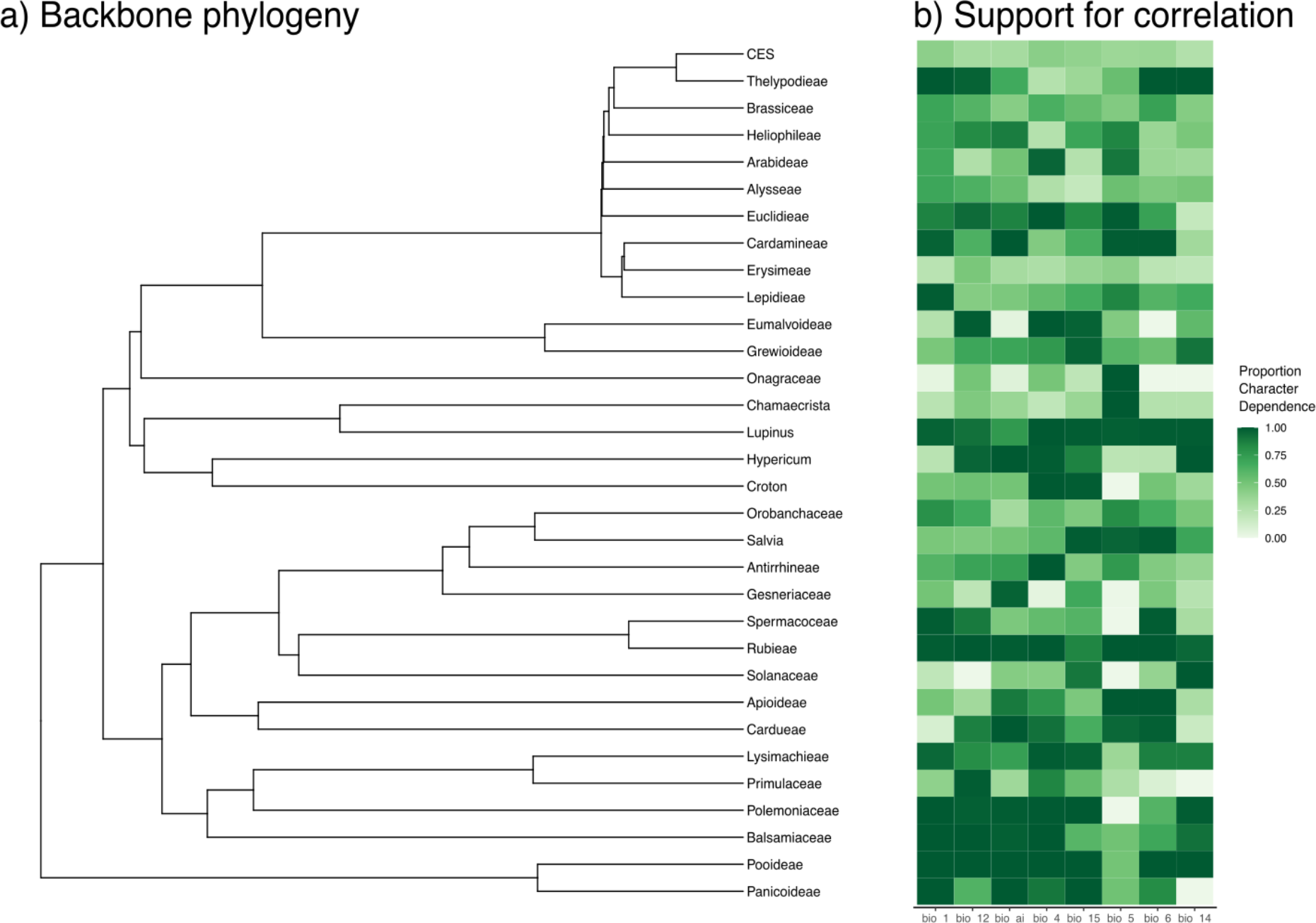
Heatmap indicating which clades have support for correlation (i.e., character dependence) for each climatic variable. The value quantifying overall model set support for orrelation is calculated as the sum of the AICc weights for the model multiplied by either 1 or CD class models, 0 for CID class models, and 0.5 for HYB models. Although HYB models are counted as 50% support correlation in this graphic, their actual interpretation will depend on specific results of the model.

## Results

### Multi-clade analysis and model selection with hOUwie

In general, we found mixed support for correlation depending on both the clade and climatic variable being analyzed (Figure 2). Certain clades, such as *Lupinus* and Pooideae, had consistent support for some form of character dependence, whereas other clades, such as Onagraceae and *Chamaecrista*, showed little correlation between life history strategy and climatic niche evolution. However, these patterns are only broad overviews and do not distinguish between character-dependent relationships with different underpinnings (i.e., both a variable-*θ* model and a variable-*σ*^2^ model are considered character-dependent). To determine whether our hypotheses were supported by the modeling results, we examined the model-averaged expected values and variances for annual and perennial lineages.

### Clade-specific parameter estimates and results

We found clade-specific differences between annuals and perennials when we examined the four climatic variables related to temperature. For BIO1 (mean annual temperature; Table S1), the difference in expected value ranged from 10.04 °C higher for annuals in Euclidieae to 4.7°C higher for perennials in Balsaminaceae. On average, the expected difference between annuals and perennials was 1.26°C warmer in annuals. All clades but Balsaminaceae, *Croton*, Erysimeae, Eumalvoideae, *Hypericum*, Onagraceae, Primulaceae, and Solanaceae had a pattern of higher expected temperature for annuals. For BIO4 (temperature seasonality; Table S2), the difference in expected temperature seasonality ranged from 4.31°C standard deviations higher for annuals in Balsaminaceae to 0.39°C standard deviations higher for perennials in Spermacoceae. On average, the expected difference between annuals and perennials was a temperature seasonality of 0.42°C standard deviations greater in annuals. All clades but Brassiceae, Gesneriaceae, Lysimachieae, Orobanchaceae, Rubieae, and Spermacoceae showed the dominant pattern of higher temperature seasonality for annuals. For BIO5 (maximum temperature of the warmest month; Table S3), the difference in expected maximum temperature ranged from 14.85°C greater for annuals in Euclidieae to 0.17°C greater for perennials in Balsaminaceae. On average, the expected difference between annuals and perennials was a maximum temperature in the warmest month of 1.81°C greater in annuals. All clades except Balsaminaceae presented this pattern. Finally, for BIO6 (minimum temperature of the coldest month; Table S4), the difference in expected minimum temperature ranged from 9.46°C colder for annuals in *Croton* to 6.79°C colder for perennials in Pooideae. On average, the expected difference between annuals and perennials was a minimum temperature of 0.98°C colder for perennials. All clades except Balsaminaceae, *Croton*, Erysimeae, Grewioideae, Lepidieae, Onagraceae, Panicoideae, Polemoniaceae, Primulaceae, and Solanaceae presented this pattern.

We also examined three climatic variables related to precipitation. For BIO12 (mean annual precipitation; Table S5), the difference in expected precipitation ranged from 198.46mm greater for annuals in Thelypodieae to 618.71mm greater for perennials in Balsaminaceae. On average, the expected difference between annuals and perennials was 63.57mm more precipitation in perennials. Clades that had greater expected annual precipitation in annuals are Brassiceae, Cardamineae, the CES clade (Brassicaceae tribes Cremolobeae, Eudemeae, Schizopetaleae), *Chamaecrista*, Gesneriaceae, Lysimachieae, Orobanchaceae, Spermacoceae, and Thelypodieae. For BIO14 (precipitation of the driest month; Table S6), the difference in expected precipitation of the driest month ranged from 1.49mm greater for annuals in Brassiceae to 28.90mm greater for perennials in *Hypericum*. On average, the expected difference between annuals and perennials was 3.66 mm more precipitation during the driest month in perennials. Clades that had greater expected precipitation during the driest month in annuals are Apioideae, Brassiceae, Cardamineae, the CES clade, *Chamaecrista*, *Croton*, Erysimeae, Orobanchaceae, *Salvia*, and Thelypodieae. For BIO15 (precipitation seasonality; Table S7), the difference in expected precipitation seasonality ranged from a coefficient of variation (the ratio of the standard deviation to the mean) of 21.27 for annuals in Grewioideae to a coefficient of variation that was 18.46 for perennials in *Croton*. On average, the coefficient of variation was 1.24 more seasonal in annuals than perennials. Clades that had greater precipitation seasonality in perennials are Antirrhineae, Apioideae, Brassiceae, Cardamineae, Cardueae, the CES clade, *Chamaecrista*, *Croton*, Erysimeae, Euclidieae, and Orobanchaceae.

Finally, for AI (Table S8), the difference in expected climatic value ranged from 0.14 higher (i.e., more humid) for annuals in Gesneriaceae to 0.34 higher in perennials for *Lupinus*. On average, the humidity was greater by 0.069AI for perennials. Clades that showed a greater climatic preference for humidity in annuals are Brassiceae, Gesneriaceae, Onagraceae, Orobanchaceae, Spermacoceae, and Thelypodieae.

### General patterns in climatic preferences

There were few consistently significant differences across several clades in terms of the expected variance (Figure 3). Although there were clade-specific differences in the evolutionary rates of climatic niche evolution for several of the climatic variables, only one showed a significant difference when accounting for all clades. Specifically, the minimum temperature of the coldest month was significantly more variable for perennials than annuals (Figure 3d; p < 0.05). This suggests higher rates of macroevolutionary change in the optimal minimum temperatures for perennial lineages in general.

**Figure 3.**
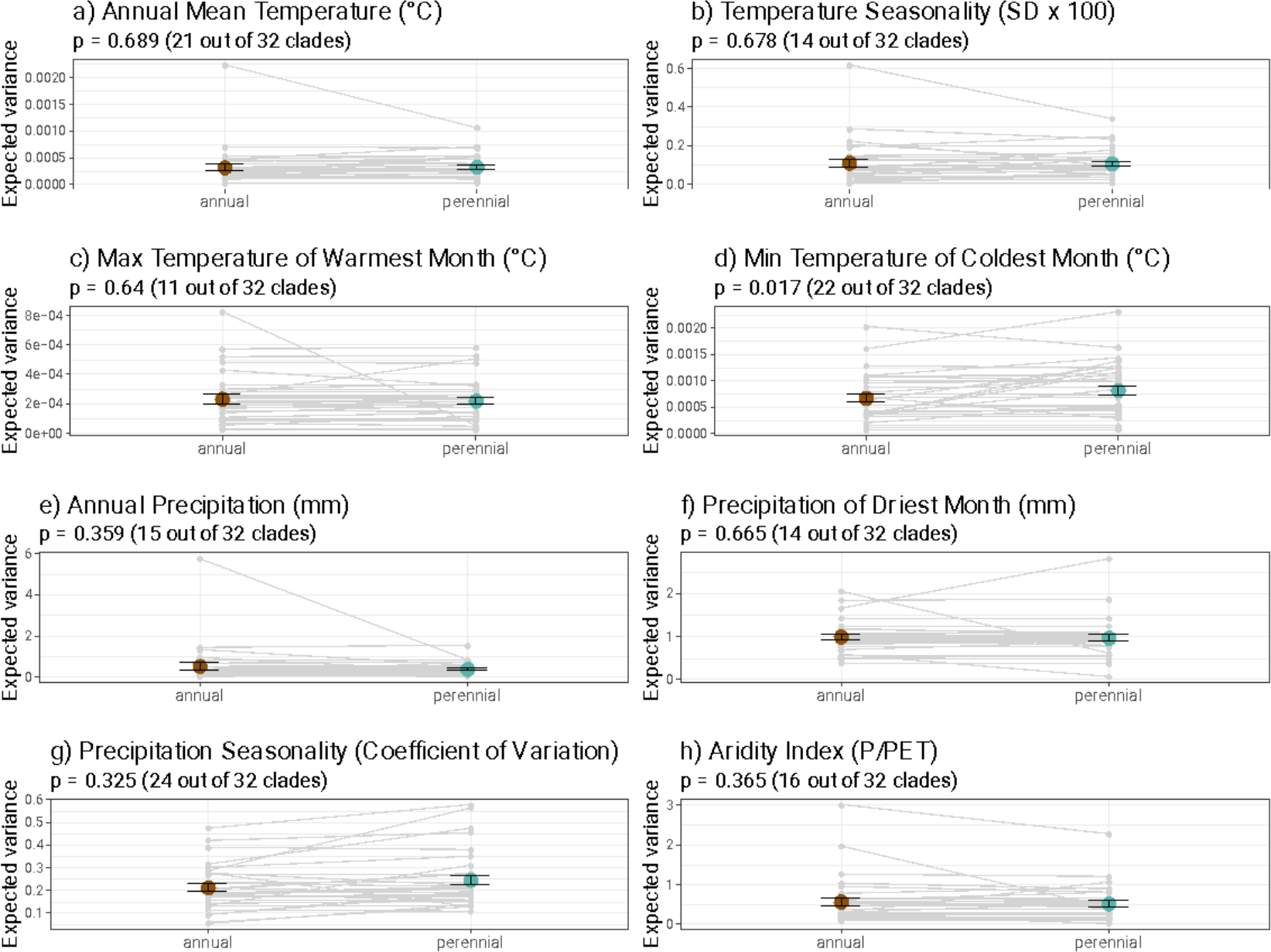
Comparison of averaged expected variance in annuals and perennials for eight climatic variables and across 32 clades. Grey lines represent individual clade comparisons between estimates associated with each observed state. Foreground points are the mean values of each expected value. p-values result from t-tests incorporating phylogenetic information.

When considering all clades, several climatic variables showed consistent differences in expected values between annual and perennial strategies (Figure 4). Mean annual temperature, maximum temperature of the warmest month, minimum temperature of the coldest month, precipitation of the driest month, and Aridity Index all showed differences at a significance value of p < 0.05 when conducting paired t-tests that incorporated phylogenetic information. In general, annuals tended to prefer warmer and drier habitats than perennials, but the most consistent pattern was that of the maximum temperature of the warmest month, in which all but one clade showed a pattern of annuals preferentially being distributed in climates prone to extreme heat (Figure 4c).

**Figure 4.**
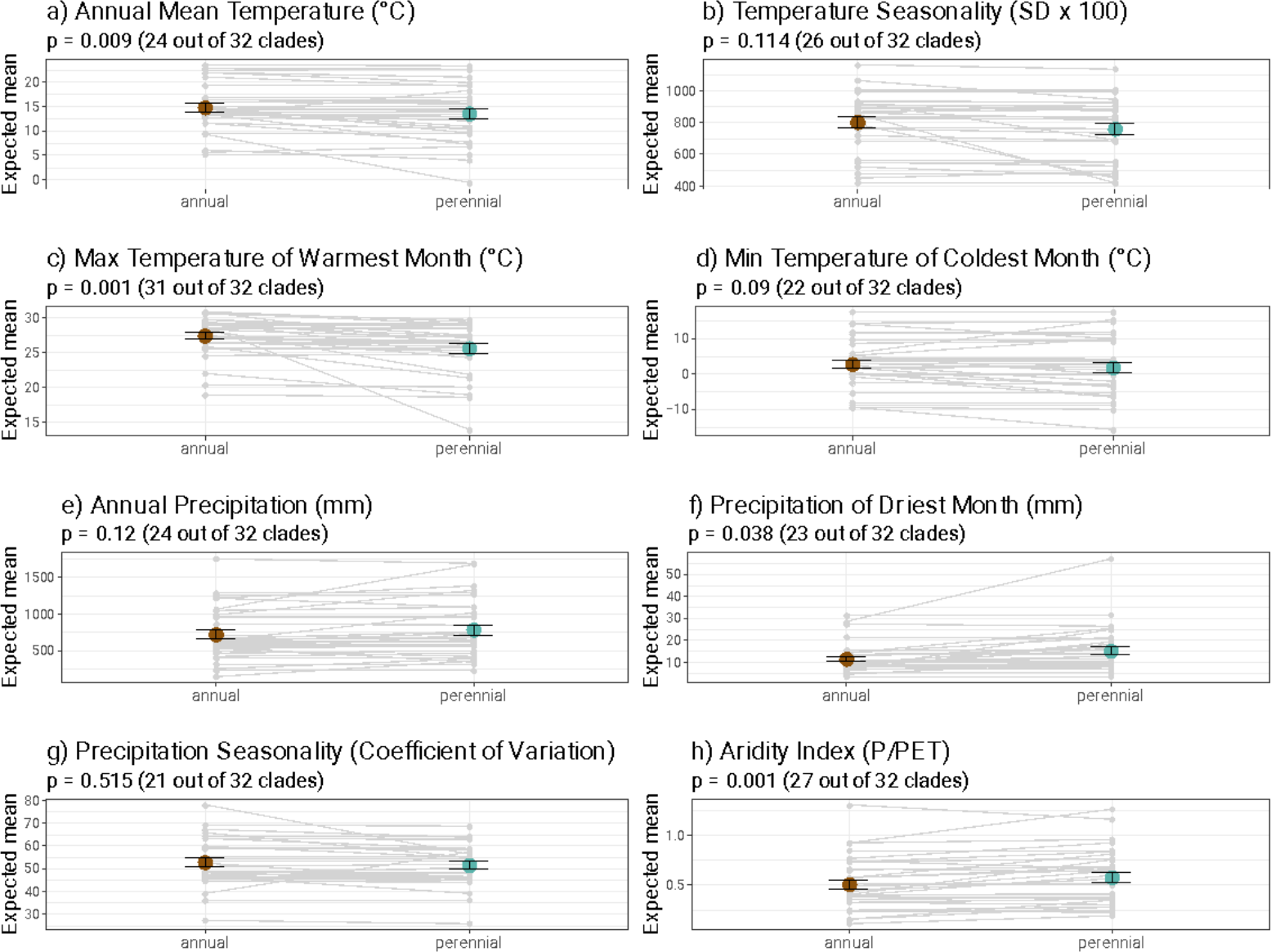
Comparison of averaged expected values in annuals and perennials for eight climatic variables and across 32 clades. Gray lines represent individual clade comparisons between estimates associated with each observed state. Foreground points are the mean values of each expected value. p-values result from phylogenetically t-tests incorporating phylogenetic information analyses.

Transition rates from annual to perennial (0.082 ± 0.50 transitions per million years) tended to be higher than from perennial to annual (0.040 ± 0.87 transitions per million years). We note that in cases where the discrete character was influenced by the continuous character (CD models), there is the potential for a great deal of variation in the ancestral state (Figure S1). This is because, even though a purely discrete process may favor an entirely annual or perennial life history when accounting for a reconstruction of the climatic niche, the most probable discrete state will also depend on the continuous character distribution. For example, the ancestral state of Primulaceae had a marginal probability of 65% annual life history when being modeled jointly with annual precipitation, but it had a marginal probability of 65% perennial life history when modeled jointly with the Aridity Index.

## Discussion

### Annual strategy as a heat-avoidance mechanism

The most consistent pattern we found across almost all analyzed clades relates to their response to extreme heat. In 31 out of the 32 clades, we found that annuals exhibit consistently higher expected values for the maximum temperature of the warmest month. This points towards a generality in the way flowering plants evolve to survive in areas subject to extreme heat, where adult mortality is high, and an optimal option may be surviving as a seed through the hottest seasons (Angert et al. 2007; Venable 2007). Both annuals and perennials are probably equally sensitive to heat stress in their adult forms (Raunkiaer 1934; Teskey et al. 2015), but an annual plant can evade the hottest season in the form of a seed, which is one of the most resistant plant structures (e.g., Janzen 1984).

In *Impatiens* (Balsaminaceae), the group that went against this general pattern, many perennials are native to the warmer tropical areas, whereas many of the annuals species occur in temperate regions of North America, Europe, and Asia (Grey-Wilson 1980; Ruchisansakun et al. 2016). They are mainly summer annuals (i.e., complete their life cycle during the summer), contrasting with other species in our dataset which are winter annuals (complete life cycle during the winter, e.g., Mulroy and Rundel 1977). Though, to our knowledge, there is no list of species on a global scale that distinguishes winter from summer annuals, we suspect that annuals consistently show higher expected values for the maximum temperature of the warmest month because most annuals in our dataset are probably winter annuals. This possibility would be consistent with the observation that Mediterranean and subtropical deserts, where the hot summers are the most unfavorable season for plants, generally favor the evolution of annuals. From an evolutionary standpoint, this further supports the lack of alternative pathways for heat tolerance in vegetative structures in plants. This is a worrying scenario for most environments dominated by perennials because extreme heat and heat waves are becoming increasingly frequent in these areas (Teskey et al. 2015).

### Annuals do not have faster rates of climatic niche evolution

Previous literature suggests that lineages with shorter generation times have faster rates of evolution (e.g., Mooers and Harvey 1994; Smith and Beaulieu 2009), but we found that this is not the case for annuals. This may be due to the fact that, although annuals do tend to have a faster development in their post-germination phase (Grime 1977; Friedman 2020), their generations are not necessarily shorter because annuals can also have relatively longer seed dormancy and can remain in the form of seeds for many years (Venable and Lawlor 1980; Nunney 2002; Kooyers 2015). In this way, their generation times can in fact be much longer in the pre-germination phase, leading to the incorrect assumption that the visible aboveground, post-germination phase represents the whole life cycle.

Another reason annuals may not have generally faster rates of evolution than perennials is that many annuals are self-compatible due to the necessity of guaranteed fertilization in a single reproductive event (Aarssen 2000). Selfing has long been considered an evolutionary dead-end in plants (Stebbins 1950) because inbreeding depression reduces the genetic diversity of selfing populations, precluding adaptation to changing environments (Takebayashi and Morrell 2001; Escobar et al. 2010; Shimizu and Tsuchimatsu 2015; but see Igic and Busch 2013). This may constrain rates of niche evolution in annuals despite their generally higher vagility. In areas of constant disturbance, such as those most influenced by human activity, annuals will be favored due to their higher vagility and their short reproductive window between germination and seed dispersal (Grime 1977). Though this may make them seem like better invaders, they may nonetheless be poor competitors compared to perennials in more stable environments. Therefore, and despite their general association with traits linked to vagility, the annual strategy may restrict plant lineages to a few types of environments where they can outcompete perennial plants – that is, regions prone to extreme heat (see *Annual strategy as a heat-avoidance mechanism*).

### Lack of general rules for most variables, including seasonality and precipitation

The accessibility to data and methods which can be used to test hypotheses about trait evolution with phylogenetic comparative methods has increased, and with that, multiple studies have found that temperature, precipitation, and seasonality variables are relevant in explaining the evolution of different life history strategies in plants (e.g., Fiz et al. 2002; Evans et al. 2005; Humphreys and Linder 2013; Ogburn and Edwards 2015; Monroe et al. 2019). Our analyses of multiple clades show that some of these previously documented patterns are not, in fact, general across flowering plants but are instead specific to certain clades or areas. For instance, we found no significant difference in the expected values for mean annual precipitation across all clades. The lack of a strong signal for this variable as an important factor in the evolution of annual strategy was unanticipated. We did recover a significant difference between expected values for precipitation of the driest month (p < 0.05) and Aridity Index (p <0.01), with annuals tending to present lower expected values, but this pattern was not observed in 8 out of 32 clades analyzed. In one-fourth of the clades, it was actually perennials that were expected to prefer drier conditions. One potential reason for this lack of strong correlation with precipitation may be the existence of other compensatory mechanisms which deal with extreme drought in perennial plants, allowing them to forgo transitions to annual life histories. Several mechanisms of vegetative tolerance to desiccation have evolved in perennials, including changes in photosynthesis pathways (Ehleringer et al. 1991), possession of subterraneous structures (Howard et al. 2019), succulence of leaves and stems (Ogburn and Edwards 2015), and senescence of photosynthesis structures during dry seasons (Munné-Bosch and Alegre 2004).

A similar lack of correlation with life history was found for all variables related to seasonality as well as for the minimum temperature of the coldest month, which is a variable associated with freezing temperatures. In these cases, optima for annuals and perennials were not significantly different from each other across all clades, meaning that there is little support for any role of these climatic variables in predictably governing life history evolution across plant clades. If these variables are related to life history evolution in these clades, the relationships are likely weak and particular to these clades’ geographical distributions. For example, in groups where species distribution varies from dry lowland to humid alpine environments, such as *Lupinus* (Drummond et al. 2012; Givnish 2015) and the Brassicaeae tribe Arabideae (Koch et al. 2012), perennials were found to have lower expected values for minimum experienced temperature. In those cases, perennials may be associated with a frost tolerance strategy due to somewhat well-distributed events of frost in mountains that lead to high seedling mortality in annuals (“winter by night and summer by day”; Givnish 2015). However, in groups such as Balsaminaceae, Onagraceae, and Solanaceae, where their distribution ranges from tropical to temperate biomes (e.g., Wagner et al. 2007) and many perennial species are restricted to humid tropical and subtropical forests where frost does not occur, the annual strategy is found in areas where occasional events of frost are present, such as Mediterranean habitats (Pescador et al. 2018). Despite the lack of generalities for these variables across the whole of flowering plants analyzed in this study, we acknowledge their probable importance in some groups.

### Multi-clade analyses shed light on both general and clade-specific patterns

Biology is scale-dependent in terms of space (McGill 2010), time (Haldane 1956), and evolutionary hierarchy (Gould 2002). Every study must make methodological choices to focus on variables of interest at the expense of other variables at different scales. For example, in comparative biology, studies examining large phylogenies usually have little power to determine specific mechanisms underpinning evolutionary patterns (Donoghue and Edwards 2019), while small-scale studies of specific clades, for various reasons, are often limited in their ability to explain broad evolutionary patterns (Beaulieu and O’Meara 2018, 2019). The multi-clade approach we used for this work, which allowed us to examine broad patterns as well as make inferences about the causes of clade-specific patterns, aims to combine the advantages of both large- and small-scale studies (Vasconcelos 2022). Due to their advantages, multi-clade studies have recently become popular in comparative biology (e.g., Mayrose et al. 2011, 2015; Vasconcelos et al. 2020, 2021; Miller et al. 2021). However, the approach has limitations of its own. The results of our study are subject to ascertainment bias due to the necessity of studying clades containing both annual and perennial species.

The importance of examining results at multiple scales before developing generalizations is underscored by comparing our findings with those of previous studies that did not use a multi-clade phylogenetic comparative approach. For example, in classical botany, annuality is generally considered to be a “derived” position in flowering plants (e.g., Stebbins 1965; Soltis et al. 2013). However, our ancestral state reconstructions recovered an annual root state with greater than 50% certainty in 13 out of 33 clades. Additionally, in some clades, the annual root state was variable, with Apioideae, Rubieae, and Balsaminaceae showing strong support for either annual or perennial root states depending on the climatic variable being modeled. This highlights both the importance of joint modeling as well as the inherent uncertainty of reconstructing ancestral states, especially because climate has been found to be an important factor influencing the evolution of many discrete plant traits, including fruit type (Vasconcelos et al. 2021) and underground storage organs (Tribble et al. 2021). Thus, the multi-clade approach allowed us to not only shed skeptical light on generalizations about transitions between annuality and perenniality, among others, but also identify clades that share evolutionary patterns as well as inspire future work to uncover possible shared mechanisms underlying those patterns. A renewed focus on the climatic specificities of individual clades will likely continue to upend long-held beliefs in plant biology, especially as new phylogenetic comparative tools like joint modeling proliferate and become more widely used.

### Conclusion

This study provides a broad-scale analysis of life history evolution in flowering plants in relation to their distribution across a climatic gradient. We show how a multi-clade analysis can challenge previous ideas which are generally based on analyses of few taxonomic groups. As predicted, we found mixed support for most climatic variables tested due to clade-specific evolutionary patterns. However, this approach also allowed us to identify at least one generality in the long-term responses of life history evolution in relation to climate. Temperature variables, and specifically extreme heat, were found to have consistent effects in almost all clades, pointing towards a possible generality in the evolution of the annual semelparous strategy as a heat avoidance mechanism, possibly due to the lack of alternative evolutionary pathways to survive heat stress in plants. Finally, we show how climatic variables have a strong influence on the evolution of correlated discrete traits when a joint modeling approach is employed.

## Acknowledgements

We thank the members of the Beaulieu and Smith labs for their comments and discussion of the ideas presented here.

## Author contributions

JDB, EH, JMB, and TV designed the study. TV collected and organized the datasets. JDB conducted the analyses. JDB, EH, JMB, and TV wrote the paper.

## Data availability

All code necessary to conduct the analyses and original tables are available at https://github.com/jboyko/life_history_houwie

## References

Aarssen LW. 2000. Why are most selfers annuals? A new hypothesis for the fitness benefit of selfing. Oikos 89: 606–612.

Andreasen K, Baldwin BG. 2001. Unequal evolutionary rates between annual and perennial lineages of checker mallows (Sidalcea, Malvaceae): evidence from 18S–26S rDNA internal and external transcribed spacers. Molecular Biology and Evolution 18: 936–944.

Angert AL, Huxman TE, Barron-Gafford GA, Gerst KL, Venable DL. 2007. Linking growth strategies to long-term population dynamics in a guild of desert annuals. Journal of Ecology 95: 321–331.

Arévalo R, van Ee BW, Riina R, Berry PE, Wiedenhoeft AC. 2017. Force of habit: shrubs, trees and contingent evolution of wood anatomical diversity using Croton (Euphorbiaceae) as a model system. Annals of Botany 119: 563–579.

Banasiak Ł, Piwczyński M, Uliński T, Downie SR, Watson MF, Shakya B, Spalik K. 2013. Dispersal patterns in space and time: a case study of Apiaceae subfamily Apioideae. Journal of Biogeography 40: 1324–1335.

Beaulieu JM, Jhwueng D-C, Boettiger C, O’Meara BC. 2012. Modeling stabilizing selection: expanding the Ornstein–Uhlenbeck model of adaptive evolution. Evolution 66: 2369–2383.

Beaulieu, J.M., O’Meara, B.C. 2016. Detecting Hidden Diversification Shifts in Models of Trait-Dependent Speciation and Extinction. Systematic Biology 65: 583–601.

Beaulieu JM, O’Meara BC. 2018. Can we build it? Yes we can, but should we use it? Assessing the quality and value of a very large phylogeny of campanulid angiosperms. American Journal of Botany 105: 417–432.

Beaulieu, JM, O’Meara BC. 2019. Diversity and skepticism are vital for comparative biology: a response to Donoghue and Edwards (2019). American Journal of Botany 106: 613–617.

Billings WD, Mooney HA. 1968. The ecology of arctic and alpine plants. Biological Reviews 43: 481–529.

Bivand RS, Pebesma EJ, Gómez-Rubio V, Pebesma EJ. 2008. Applied spatial data analysis with R. New York, NY: Springer.

Boyko JD, Beaulieu JM. 2022. Reducing the biases in false correlations between discrete characters. Systematic Biology. doi: https://doi.org/10.1093/sysbio/syac066.

Boyko JD, O’Meara BC, Beaulieu JM. 2022. Jointly modeling the evolution of discrete and continuous traits. EcoEvoRxiv. doi: 10.32942/osf.io/fb8k7.

Brummitt RK, Pando F, Hollis S, Brummitt NA. 2001. Plant taxonomic database standards no. 2. World geographical scheme for recording plant distributions, 2^nd^ edn. Pittsburgh, PA: Published for the International Working Group on Taxonomic Databases For Plant Sciences (TDWG) by the Hunt Institute for Botanical Documentation, Carnegie Mellon University.

Butler MA, King AA. 2004. Phylogenetic comparative analysis: a modeling approach for adaptive evolution. The American Naturalist 164: 683–695.

Caetano, D.S., O’Meara, B.C., Beaulieu, J.M. 2018. Hidden state models improve state-dependent diversification approaches, including biogeographical models: HMM and the adequacy of SSE models. Evolution 72: 2308–2324.

Chamberlain SA, Boettiger C. 2017. R Python, and Ruby clients for GBIF species occurrence data. PeerJ Preprints 5: e3304v1.

Chamberlain SA, Szöcs E. 2013. taxize: taxonomic search and retrieval in R. F1000Research 2: 191.

Cole LC. 1954. The population consequences of life history phenomena. The Quarterly Review of Biology 29: 103–137.

Donoghue MJ, Edwards EJ. 2019. Model clades are vital for comparative biology, and ascertainment bias is not a problem in practice: a response to Beaulieu and O’Meara (2018). American Journal of Botany 106: 327–330.

Drummond CS, Eastwood RJ, Miotto STS, Hughes CE. 2012. Multiple continental radiations and correlates of diversification in Lupinus (Leguminosae): testing for key innovation with incomplete taxon sampling. Systematic Biology 61: 443–460.

Edwards EJ, Chatelet DS, Chen B-C, Ong JY, Tagane S, Kanemitsu H, Tagawa K, Teramoto K, Park B, Chung K-F, Hu J-M, Yahara T, Donoghue MJ. 2017. Convergence, consilience, and the evolution of temperate deciduous forests. The American Naturalist 190: S87–S104.

Ehleringer JR, Sage RF, Flanagan LB, Pearcy RW. 1991. Climate change and the evolution of C4 photosynthesis. Trends in Ecology & Evolution 6: 95–99.

Ehrendorfer F, Barfuss MHJ, Manen J-F, Schneeweiss GM. 2018. Phylogeny, character evolution and spatiotemporal diversification of the species-rich and world-wide distributed tribe Rubieae (Rubiaceae). PLoS ONE 13: e0207615.

Escobar JS, Cenci A, Bolognini J, Haudry A, Laurent S, David J, Glémin S. 2010. An integrative test of the dead-end hypothesis of selfing evolution in Triticeae (Poaceae). Evolution 64: 2855–2872.

Evans MEK, Hearn DJ, Hahn WJ, Spangle JM, Venable DL. 2005. Climate and life-history evolution in evening primroses (Oenothera, Onagraceae): A phylogenetic comparative analysis. Evolution 59: 1914–1927.

Fiz O, Valcárcel V, Vargas P. 2002. Phylogenetic position of Mediterranean Astereae and character evolution of daisies (Bellis, Asteraceae) inferred from nrDNA ITS sequences. Molecular Phylogenetics and Evolution 25: 157–171.

Freyman WA, Höhna S. 2019. Stochastic character mapping of state-dependent diversification reveals the tempo of evolutionary decline in self-compatible Onagraceae lineages. Systematic Biology 68: 505–519.

Friedman J. 2020. The evolution of annual and perennial plant life histories: ecological correlates and genetic mechanisms. Annual Review of Ecology, Evolution, and Systematics 51: 461–481.

Givnish TJ. 2015. Adaptive radiation versus ‘radiation’ and ‘explosive diversification’: why conceptual distinctions are fundamental to understanding evolution. New Phytologist 207: 297–303.

Gorospe JM, Monjas D, Fernández-Mazuecos M. 2020. Out of the Mediterranean region: worldwide biogeography of snapdragons and relatives (tribe Antirrhineae, Plantaginaceae). Journal of Biogeography 47: 2442–2456.

Gould SJ 2002. The structure of evolutionary theory. Cambridge, MA: Harvard University Press.

Grey-Wilson C. 1980. Impatiens of Africa. Boca Raton, FL: CRC Press.

Grime JP. 1977. Evidence for the existence of three primary strategies in plants and its relevance to ecological and evolutionary theory. The American Naturalist 111: 1169–1194.

Haldane JBS. 1956. Time in biology. Science Progress 154: 385–402.

Hansen TF. 1997. Stabilizing selection and the comparative analysis of adaptation. Evolution 51: 1341–1351.

Hijmans RJ, Van Etten J, Cheng J, Mattiuzzi M, Sumner M, Greenberg JA, Lamigueiro OP, Bevan A, Racine EB, Shortridge A. 2015. raster: geographic data analysis and modeling. R package version 3.6-3.

Hoorn C, van der Ham R, de la Parra F, Salamanca S, ter Steege H, Banks H, Star W, van Heuven BJ, Langelaan R, Carvalho FA, Rodriguez-Forero G, Lagomarsino LP. 2019. Going north and south: the biogeographic history of two Malvaceae in the wake of Neogene Andean uplift and connectivity between the Americas. Review of Palaeobotany and Palynology 264: 90–109.

Howard CC, Folk RA, Beaulieu JM, Cellinese N. 2019. The monocotyledonous underground: global climatic and phylogenetic patterns of geophyte diversity. American Journal of Botany 106: 850–863.

Huang X-C, German DA, Koch MA. 2020. Temporal patterns of diversification in Brassicaceae demonstrate decoupling of rate shifts and mesopolyploidization events. Annals of Botany 125: 29–47.

Humphreys AM, Linder HP. 2013. Evidence for recent evolution of cold tolerance in grasses suggests current distribution is not limited by (low) temperature. New Phytologist 198: 1261–1273.

Igic B, Busch JW. 2013. Is self-fertilization an evolutionary dead end? New Phytologist 198: 386–397.

Janzen DH. 1984. Dispersal of small seeds by big herbivores: foliage is the fruit. The American Naturalist 123: 338–353.

Karger DN, Conrad O, Böhner J, Kawohl T, Kreft H, Soria-Auza RW, Zimmermann NE, Linder HP, Kessler M. 2017. Climatologies at high resolution for the earth’s land surface areas. Scientific Data 4: 1–20.

Koch MA, Karl R, German DA, Al-Shehbaz IA. 2012. Systematics, taxonomy and biogeography of three new Asian genera of Brassicaceae tribe Arabideae: an ancient distribution circle around the Asian high mountains. TAXON 61: 955–969.

Kooyers NJ. 2015. The evolution of drought escape and avoidance in natural herbaceous populations. Plant Science 234: 155–162.

Kriebel R, Drew B, González-Gallegos JG, Celep F, Heeg L, Mahdjoub MM, Sytsma KJ. 2020. Pollinator shifts, contingent evolution, and evolutionary constraint drive floral disparity in Salvia (Lamiaceae): evidence from morphometrics and phylogenetic comparative methods. Evolution 74: 1335–1355.

Labra A, Pienaar J, Hansen TF. 2009. Evolution of thermal physiology in Liolaemus lizards: adaptation, phylogenetic inertia, and niche tracking. The American Naturalist 174: 204–220.

Linder HP, Lehmann CER, Archibald S, Osborne CP, Richardson DM. 2018. Global grass (Poaceae) success underpinned by traits facilitating colonization, persistence and habitat transformation. Biological Reviews 93: 1125–1144.

Mayrose I, Zhan SH, Rothfels CJ, Arrigo N, Barker MS, Rieseberg LH, Otto SP. 2015. Methods for studying polyploid diversification and the dead end hypothesis: a reply to Soltis et al. (2014). New Phytologist 206: 27–35.

Mayrose I, Zhan SH, Rothfels CJ, Magnuson-Ford K, Barker MS, Rieseberg LH, Otto SP. 2011. Recently formed polyploid plants diversify at lower rates. Science 333: 1257.

McGill BJ. 2010. Matters of scale. Science 328: 575–576.

Miller EC, Mesnick SL, Wiens JJ. 2021. Sexual dichromatism is decoupled from diversification over deep time in fishes. The American Naturalist 198: 232–252.

Monroe JG, Gill B, Turner KG, McKay JK. 2019. Drought regimens predict life history strategies in Heliophila. New Phytologist 223: 2054–2062.

Mooers AØ, Harvey PH. 1994. Metabolic rate, generation time, and the rate of molecular evolution in birds. Molecular Phylogenetics and Evolution 3: 344–350.

Mulroy TW, Rundel PW. 1977. Annual plants: adaptations to desert environments. BioScience 27: 109–114.

Munné-Bosch S, Alegre L. 2004. Die and let live: leaf senescence contributes to plant survival under drought stress. Functional Plant Biology 31: 203–216.

Neupane S, Lewis PO, Dessein S, Shanks H, Paudyal S, Lens F. 2017. Evolution of woody life form on tropical mountains in the tribe Spermacoceae (Rubiaceae). American Journal of Botany 104: 419–438.

Nunney L. 2002. The effective size of annual plant populations: the interaction of a seed bank with fluctuating population size in maintaining genetic variation. The American Naturalist 160: 195–204.

Nürk N, Scheriau C, Madriñán S. 2013. Explosive radiation in high Andean Hypericum— rates of diversification among New World lineages. Frontiers in Genetics 4: 175.

Ogburn MR, Edwards EJ. 2015. Life history lability underlies rapid climate niche evolution in the angiosperm clade Montiaceae. Molecular Phylogenetics and Evolution 92: 181–192.

Pannell JR, Auld JR, Brandvain Y, Burd M, Busch JW, Cheptou P-O, Conner JK, Goldberg EE, Grant A-G, Grossenbacher DL, Hovick SM, Igic B, Kalisz S, Petanidou T, Randle AM, de Casas RR, Pauw A, Vamosi JC, Winn AA. 2015. The scope of Baker’s law. New Phytologist 208: 656–667.

Paradis E, Claude J, Strimmer K. 2004. APE: analyses of phylogenetics and evolution in R language. Bioinformatics 20: 289–290.

Park DS, Potter D. 2015. Why close relatives make bad neighbours: phylogenetic conservatism in niche preferences and dispersal disproves Darwin’s naturalization hypothesis in the thistle tribe. Molecular Ecology 24: 3181–3193.

Pescador DS, Sánchez AM, Luzuriaga AL, Sierra-Almeida A, Escudero A. 2018. Winter is coming: plant freezing resistance as a key functional trait for the assembly of annual Mediterranean communities. Annals of Botany 121: 335–344.

POWO. 2022. Plants of the World Online. Facilitated by the Royal Botanic Gardens, Kew. [WWW document] URL http://www.plantsoftheworldonline.org/ [accessed 08/14/22].

Rando JG, Zuntini AR, Conceição AS, van den Berg C, Pirani JR, de Queiroz LP. 2016. Phylogeny of Chamaecrista ser. Coriaceae (Leguminosae) unveils a lineage recently diversified in Brazilian Campo Rupestre vegetation. International Journal of Plant Sciences 177: 3–17.

Raunkiaer C. 1934. The life forms of plants and statistical plant geography; being the collected papers of C. Raunkiaer. Oxford: Clarendon Press.

Revell L.J. 2012. phytools: an R package for phylogenetic comparative biology (and other things). Methods in Ecology and Evolution 3: 217–223.

Ricklefs RE, Renner SS. 1994. Species richness within families of flowering plants. Evolution 48: 1619–1636.

Roalson EH, Roberts WR. 2016. Distinct processes drive diversification in different clades of Gesneriaceae. Systematic Biology 65: 662–684.

Rose JP, Kleist TJ, Löfstrand SD, Drew BT, Schönenberger J, Sytsma KJ. 2018. Phylogeny, historical biogeography, and diversification of angiosperm order Ericales suggest ancient Neotropical and East Asian connections. Molecular Phylogenetics and Evolution 122: 59–79.

Ruchisansakun S, van der Niet T, Janssens SB, Triboun P, Techaprasan J, Jenjittikul T, Suksathan P. 2016. Phylogenetic analyses of molecular data and reconstruction of morphological character evolution in Asian Impatiens section Semeiocardium (Balsaminaceae). Systematic Botany 40: 1063–1074.

Salariato DL, Zuloaga FO, Franzke A, Mummenhoff K, Al-Shehbaz IA. 2016. Diversification patterns in the CES clade (Brassicaceae tribes Cremolobeae, Eudemeae, Schizopetaleae) in Andean South America. Botanical Journal of the Linnean Society 181: 543–566.

Särkinen T, Bohs L, Olmstead RG, Knapp S. 2013. A phylogenetic framework for evolutionary study of the nightshades (Solanaceae): a dated 1000-tip tree. BMC Evolutionary Biology 13: 214.

Schliep KP. 2011. phangorn: phylogenetic analysis in R. Bioinformatics 27: 592–593.

Schneider AC, Moore AJ. 2017. Parallel Pleistocene amphitropical disjunctions of a parasitic plant and its host. American Journal of Botany 104: 1745–1755.

Shimizu KK, Tsuchimatsu T. 2015. Evolution of selfing: recurrent patterns in molecular adaptation. Annual Review of Ecology, Evolution, and Systematics 46: 593–622.

Smith SA, Beaulieu JM. 2009. Life history influences rates of climatic niche evolution in flowering plants. Proceedings of the Royal Society B: Biological Sciences 276: 4345–4352.

Smith SA, Brown JW. 2018. Constructing a broadly inclusive seed plant phylogeny. American Journal of Botany 105: 302–314.

Soltis DE, Mort ME, Latvis M, Mavrodiev EV, O’Meara BC, Soltis PS, Burleigh JG, de Casas RR. 2013. Phylogenetic relationships and character evolution analysis of Saxifragales using a supermatrix approach. American Journal of Botany 100: 916–929.

Spriggs EL, Christin P-A, Edwards EJ. 2014. C4 photosynthesis promoted species diversification during the Miocene grassland expansion. PLoS ONE 9: e97722.

Stearns SC. 1992. The evolution of life histories. Oxford, UK: Oxford University Press.

Stebbins GL. 1950. Variation and evolution in plants. New York, NY: Columbia University Press.

Stebbins, GL. 1965. The probable growth habits of the earliest flowering plants. Annals of the Missouri Botanical Garden 52: 457–468.

Stebbins GL. 1974. Flowering plants: evolution above the species level. Cambridge, MA: Belknap Press of Harvard University Press.

Takebayashi N., Morrell PL. 2001. Is self-fertilization an evolutionary dead end? Revisiting an old hypothesis with genetic theories and a macroevolutionary approach. American Journal of Botany 88: 1143–1150.

Teskey R, Wertin T, Bauweraerts I, Ameye M, Mcguire MA, Steppe K. 2015. Responses of tree species to heat waves and extreme heat events. Plant, Cell & Environment 38: 1699–1712.

Trabucco A, Zomer RJ. 2019. Global Aridity Index and potential evapotranspiration (ET0) Climate Database v2 figshare. CGIAR consortium for spatial information. Figshare. doi: 10.6084/m9.figshare.7504448.v3.

Tribble CM, May MR, Jackson-Gain A, Zenil-Ferguson R, Specht CD, Rothfels CJ. 2021. Unearthing modes of climatic adaptation in underground storage organs across Liliales. bioRxiv. doi: 10.1101/2021.09.03.458928.

Vasconcelos T, Boyko JD, Beaulieu JM. 2021. Linking mode of seed dispersal and climatic niche evolution in flowering plants. Journal of Biogeography. doi: 10.1111/jbi.14292.

Vasconcelos TNC, Alcantara S, Andrino CO, Forest F, Reginato M, Simon MF, Pirani JR. 2020. Fast diversification through a mosaic of evolutionary histories characterizes the endemic flora of ancient Neotropical mountains. Proceedings of the Royal Society B: Biological Sciences 287: 20192933.

Vasconcelos, TNC. 2022. Discovering the Rules of Plant Biogeography Using a Trait-based Approach. EcoEvoRxiv. doi: ecoevorxiv.org/azytc.

Venable DL. 2007. Bet hedging in a guild of desert annuals. Ecology 88: 1086–1090.

Venable DL, Lawlor L. 1980. Delayed germination and dispersal in desert annuals: escape in space and time. Oecologia 46: 272–282.

de Vos JM, Hughes CE, Schneeweiss GM, Moore BR, Conti E. 2014. Heterostyly accelerates diversification via reduced extinction in primroses. Proceedings of the Royal Society B: Biological Sciences 281: 20140075.

Wagner WL, Hoch PC, Raven PH. 2007. Revised classification of the Onagraceae. Systematic Botany Monographs 83: 1–240.

Yan H-F, Zhang C-Y, Anderberg AA, Hao G, Ge X-J, Wiens JJ. 2018. What explains high plant richness in East Asia? Time and diversification in the tribe Lysimachieae (Primulaceae). New Phytologist 219: 436–448.

Zanne, A.E., Tank, D.C., Cornwell, W.K., Eastman, J.M., Smith, S.A., FitzJohn, R.G., McGlinn, D.J., O’Meara, B.C., Moles, A.T., Reich, P.B. 2014. Three keys to the radiation of angiosperms into freezing environments. Nature 506: 89–92.

