## Supplemental tables and files for "Long-term responses of life-history strategies to climatic variability in flowering plants"

***New Phytologist* Supporting Information**

Article acceptance date: N/A

The following Supporting Information is available for this article:

**Figure S1.** Mean probability of an annual root state for each of the 32 analyzed clades. Error bars show the range of probabilities for ancestral state reconstructed in the root of each phylogeny depending on a given bioclimatic variable.

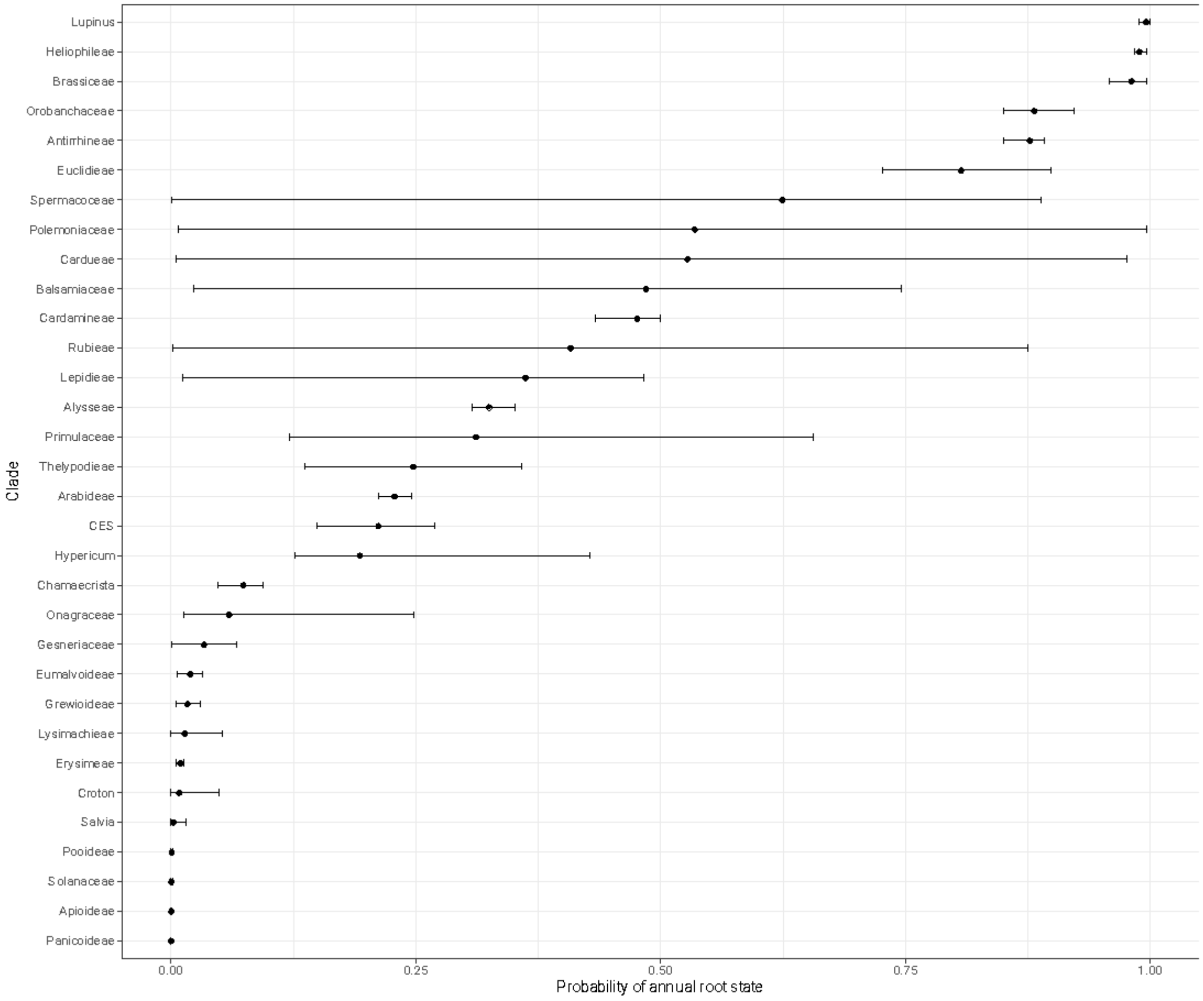

**Table S1** Parameter estimates from the model averaged hOUwie fits for BIO1

| Group.1 | Group.2 | rates | alpha | sigma.sq | theta | expected_mean | expected_var |
|---|---|---|---|---|---|---|---|
| Alysseae | annual | 0.04222935 | 6.77642682 | 0.00120016 | 11.6358211 | 11.599203 | 0.00010195 |
| Antirrhineae | annual | 0.03352804 | 0.05283134 | 2.79E-05 | 16.4969491 | 16.474252 | 0.00026284 |
| Apioideae | annual | 0.01257536 | 0.09463659 | 2.60E-05 | 17.2854379 | 13.9009453 | 0.00020198 |
| Arabideae | annual | 0.27432857 | 1.43675206 | 0.00151969 | 6.86915524 | 5.9892773 | 0.00052886 |
| Balsamiaceae | annual | 0.01477442 | 0.12344513 | 5.84E-05 | 13.0946964 | 13.4773253 | 0.00023675 |
| Brassiceae | annual | 0.05602161 | 7.38562273 | 0.0029488 | 15.075985 | 15.0753257 | 0.00019591 |
| Cardamineae | annual | 0.02703944 | 1.56747447 | 0.00061663 | 13.7721924 | 11.5653176 | 0.00042866 |
| Cardueae | annual | 0.05775609 | 0.04988559 | 3.19E-05 | 59.6526461 | 12.6869729 | 0.00032851 |
| CES | annual | 0.09746567 | 8.01E-05 | 2.26E-05 | 1494.21649 | 5.18294859 | 0.00045133 |
| Chamaecrista | annual | 0.0798664 | 0.19370296 | 1.06E-05 | 23.5948827 | 23.4885925 | 2.89E-05 |
| Croton | annual | 0.01352555 | 0.09443209 | 5.06E-05 | 22.2632167 | 22.4232046 | 0.0001998 |
| Erysimeae | annual | 0.18350651 | 0.57528604 | 0.00021464 | 9.48554426 | 9.55440502 | 0.00023344 |
| Euclidieae | annual | 0.04170806 | 1.44339278 | 0.00194283 | 9.5400535 | 9.31514576 | 0.00069451 |
| Eumalvoideae | annual | 0.04097997 | 0.07407 | 4.08E-05 | 19.3896529 | 19.3014422 | 0.00027518 |
| Gesneriaceae | annual | 0.02339613 | 0.01142421 | 5.37E-06 | 293.347346 | 21.0624018 | 0.00031562 |
| Grewioideae | annual | 0.0181161 | 0.02400397 | 4.95E-06 | 22.8486457 | 22.7632131 | 0.00010369 |
| Heliophileae | annual | 0.03608735 | 5.34179125 | 0.00021168 | 16.4494834 | 16.4494831 | 1.94E-05 |
| Hypericum | annual | 0.08462825 | 0.38829114 | 0.00015669 | 15.0683909 | 15.0818612 | 0.00020344 |
| Lepidieae | annual | 0.28809948 | 0.07966505 | 0.0006015 | 14.361843 | 14.3617829 | 0.00222598 |
| Lupinus | annual | 0.08945093 | 0.00771825 | 1.40E-05 | 13.8705263 | 13.8615744 | 0.00019324 |
| Lysimachieae | annual | 0.031292 | 0.07225652 | 6.49E-06 | 17.9111068 | 13.8900604 | 0.00016277 |
| Onagraceae | annual | 0.08565747 | 0.33838161 | 0.00030812 | 14.4855368 | 14.4851097 | 0.00047495 |
| Orobanchaceae | annual | 0.05004705 | 0.0930598 | 7.61E-05 | 13.1565371 | 13.1141786 | 0.00045983 |
| Panicoideae | annual | 0.02674436 | 5.12319916 | 0.00143838 | 21.9289641 | 21.9190316 | 0.00014045 |
| Polemoniaceae | annual | 0.01113531 | 0.0500847 | 2.15E-05 | 13.509091 | 13.5019742 | 0.00022586 |
| Pooideae | annual | 0.01881985 | 2.24398818 | 0.00058969 | 13.7395079 | 13.3865246 | 0.00014158 |
| Primulaceae | annual | 0.02811619 | 0.1964767 | 0.00015072 | 5.20605779 | 5.66541944 | 0.00038335 |
| Rubieae | annual | 0.20261902 | 14.7219208 | 0.00248467 | 14.2412054 | 14.2410898 | 8.60E-05 |
| Salvia | annual | 0.01835306 | 0.09468796 | 2.79E-05 | 16.9541833 | 15.9886691 | 0.00014706 |
| Solanaceae | annual | 0.04090887 | 0.11920195 | 7.30E-05 | 16.7514343 | 16.8145424 | 0.00030601 |
| Spermacoceae | annual | 0.03367712 | 0.10978934 | 3.28E-05 | 22.276896 | 22.0518739 | 0.00014929 |
| Thelypodieae | annual | 0.03081334 | 0.60214063 | 0.00018454 | 16.0302033 | 15.1078576 | 0.00014456 |
| Alysseae | perennial | 0.02199708 | 6.77642682 | 0.00172702 | 10.5103978 | 10.5140164 | 0.00011952 |
| Antirrhineae | perennial | 0.0120018 | 0.05283134 | 2.51E-05 | 15.7493571 | 16.1769452 | 0.00024855 |
| Apioideae | perennial | 0.01301074 | 0.09463659 | 6.74E-05 | 10.437412 | 10.4937058 | 0.00035442 |
| Arabideae | perennial | 0.07987928 | 1.43675206 | 0.00151978 | 3.9353687 | 3.94547408 | 0.0005289 |
| Balsamiaceae | perennial | 0.01437154 | 0.12344513 | 5.84E-05 | 19.2453076 | 18.2078246 | 0.00023695 |
| Brassiceae | perennial | 0.01854014 | 7.38562273 | 0.0027768 | 13.9219199 | 13.9410979 | 0.00019018 |
| Cardamineae | perennial | 0.03890428 | 1.56747447 | 0.00159958 | 7.4862357 | 7.55988708 | 0.00046145 |
| Cardueae | perennial | 0.02756332 | 0.04988559 | 3.24E-05 | 12.2762565 | 12.347128 | 0.00033127 |
| CES | perennial | 0.02950397 | 8.01E-05 | 1.17E-05 | 5.05167277 | 5.06660478 | 0.00037914 |
| Chamaecrista | perennial | 0.01107084 | 0.19370296 | 1.11E-05 | 23.3225387 | 23.327161 | 2.90E-05 |
| Croton | perennial | 0.00522272 | 0.09443209 | 1.77E-05 | 22.5238456 | 22.5227251 | 9.53E-05 |
| Erysimeae | perennial | 0.02713203 | 0.57528604 | 0.00021566 | 9.59452891 | 9.5930655 | 0.00023058 |
| Euclidieae | perennial | 0.0541651 | 1.44339278 | 0.00119481 | -1.2958894 | -0.7259879 | 0.00066824 |
| Eumalvoideae | perennial | 0.01231403 | 0.07407 | 4.02E-05 | 19.3179258 | 19.3162748 | 0.00027108 |
| Gesneriaceae | perennial | 0.00224955 | 0.01142421 | 8.21E-06 | 35.6042325 | 19.204447 | 0.00034181 |
| Grewioideae | perennial | 0.00193145 | 0.02400397 | 5.42E-06 | 22.5575966 | 22.576185 | 0.00011109 |
| Heliophileae | perennial | 0.02447892 | 5.34179125 | 0.00021505 | 15.7401801 | 15.7456377 | 1.97E-05 |
| Hypericum | perennial | 0.01696327 | 0.38829114 | 0.00015788 | 15.1333481 | 15.1298741 | 0.00020352 |
| Lepidieae | perennial | 0.24003825 | 0.07966505 | 8.02E-05 | 14.3604909 | 14.3605583 | 0.00105854 |
| Lupinus | perennial | 0.17200849 | 0.00771825 | 9.51E-05 | 13.7664455 | 13.7887147 | 0.00051338 |
| Lysimachieae | perennial | 0.02562137 | 0.07225652 | 4.35E-05 | 12.0365717 | 12.1366828 | 0.00029724 |
| Onagraceae | perennial | 0.01231289 | 0.33838161 | 0.00035963 | 14.5595015 | 14.5592252 | 0.00052732 |
| Orobanchaceae | perennial | 0.04045963 | 0.0930598 | 0.00013524 | 12.8453955 | 12.9271392 | 0.00070211 |
| Panicoideae | perennial | 0.04224172 | 5.12319916 | 0.00143819 | 20.9175867 | 20.9224078 | 0.00014036 |
| Polemoniaceae | perennial | 0.01061253 | 0.0500847 | 6.56E-05 | 13.3968991 | 13.4439605 | 0.00053359 |
| Pooideae | perennial | 0.0305447 | 2.24398818 | 0.00230108 | 7.20217553 | 7.22005428 | 0.00051208 |
| Primulaceae | perennial | 0.01059084 | 0.1964767 | 0.00015072 | 6.8216377 | 6.78214719 | 0.00038336 |
| Rubieae | perennial | 0.07609358 | 14.7219208 | 0.00645165 | 10.7557004 | 10.7626708 | 0.00021825 |
| Salvia | perennial | 0.01337889 | 0.09468796 | 2.79E-05 | 15.5519714 | 15.5541254 | 0.00014708 |
| Solanaceae | perennial | 0.01390362 | 0.11920195 | 7.30E-05 | 16.8991125 | 16.8973386 | 0.00030601 |
| Spermacoceae | perennial | 0.01348743 | 0.10978934 | 3.28E-05 | 18.8112359 | 19.747762 | 0.00014929 |
| Thelypodieae | perennial | 0.07954879 | 0.60214063 | 0.00017023 | 9.30177869 | 9.40325401 | 0.00014408 |

**Table S2** Parameter estimates from the model-averaged hOUwie fits for BIO4

| Group.1 | Group.2 | rates | alpha | sigma.sq | theta | expected_mean | expected_var |
|---|---|---|---|---|---|---|---|
| Alysseae | annual | 0.04623064 | 0.29525668 | 0.01411517 | 1001.04014 | 1000.51526 | 0.02446976 |
| Antirrhineae | annual | 0.03545918 | 0.03345322 | 0.00156237 | 838.583326 | 839.489812 | 0.02628025 |
| Apioideae | annual | 0.02391669 | 0.07987379 | 0.01081229 | 943.663154 | 933.266706 | 0.07501499 |
| Arabideae | annual | 1.1513118 | 0.68015068 | 0.02881815 | 993.317362 | 992.392174 | 0.04742367 |
| Balsamiaceae | annual | 0.0145933 | 0.00380646 | 0.00582319 | 346378.375 | 849.973796 | 0.22211988 |
| Brassiceae | annual | 0.05688646 | 5.02214681 | 0.35297626 | 865.438844 | 865.480434 | 0.03502791 |
| Cardamineae | annual | 0.03194164 | 0.3267593 | 0.09297251 | 910.867313 | 908.066406 | 0.14241586 |
| Cardueae | annual | 0.07035134 | 0.09020761 | 0.00787459 | 938.278821 | 930.337622 | 0.04564716 |
| CES | annual | 0.10411611 | 0.00666191 | 0.00365852 | 1613.95006 | 832.713664 | 0.14640274 |
| Chamaecrista | annual | 0.06874797 | 0.13331923 | 0.00168011 | 421.584619 | 419.653079 | 0.00632869 |
| Croton | annual | 0.03179358 | 0.13380144 | 0.02210604 | 944.867343 | 786.431857 | 0.07999859 |
| Erysimeae | annual | 0.1982503 | 0.05325203 | 0.01549671 | 1008.16369 | 1007.71817 | 0.12207182 |
| Euclidieae | annual | 0.04590691 | 0.57957188 | 0.00504498 | 1159.92731 | 1159.7501 | 0.00417278 |
| Eumalvoideae | annual | 0.04196715 | 0.07993348 | 0.02366537 | 573.698439 | 564.939218 | 0.13420067 |
| Gesneriaceae | annual | 0.01034297 | 0.00011715 | 0.00273603 | 479.559184 | 479.624543 | 0.19744503 |
| Grewioideae | annual | 0.01839739 | 0.05615118 | 0.00826692 | 531.339029 | 521.780875 | 0.07368693 |
| Heliophileae | annual | 0.03394285 | 0.22994972 | 0.00241149 | 680.822714 | 680.805003 | 0.0066169 |
| Hypericum | annual | 0.10949778 | 0.02634112 | 0.0587529 | 1059.13797 | 892.645913 | 0.61556299 |
| Lepidieae | annual | 0.26785015 | 1.20419534 | 0.18476812 | 867.865367 | 860.580185 | 0.07780927 |
| Lupinus | annual | 0.03317324 | 0.00150559 | 0.00411432 | 767.461383 | 766.211687 | 0.07422281 |
| Lysimachieae | annual | 0.0339976 | 0.00206873 | 6.37E-05 | 815.378812 | 825.214515 | 0.19087424 |
| Onagraceae | annual | 0.07402165 | 0.05566329 | 0.0367747 | 716.509791 | 717.691354 | 0.2833446 |
| Orobanchaceae | annual | 0.04951053 | 0.10805047 | 0.01816022 | 864.614561 | 867.539143 | 0.08662163 |
| Panicoideae | annual | 0.04060338 | 5.82108862 | 0.98246451 | 549.640242 | 549.634825 | 0.08443686 |
| Polemoniaceae | annual | 0.01199421 | 0.03094406 | 0.00421092 | 936.138183 | 924.871046 | 0.06812361 |
| Pooideae | annual | 0.02256814 | 0.59969435 | 0.05025966 | 925.96754 | 916.432884 | 0.05014514 |
| Primulaceae | annual | 0.03228184 | 2.55118575 | 0.19729472 | 1067.51702 | 1065.14705 | 0.03996562 |
| Rubieae | annual | 0.13120271 | 0.95176001 | 0.01740108 | 885.231527 | 885.289456 | 0.00988396 |
| Salvia | annual | 0.03575222 | 0.04632607 | 0.01112849 | 889.904718 | 756.566801 | 0.12203424 |
| Solanaceae | annual | 0.03324896 | 0.10759233 | 0.0275607 | 565.977753 | 547.629476 | 0.12826102 |
| Spermacoceae | annual | 0.05419206 | 0.0574132 | 0.04481367 | 426.02468 | 451.752564 | 0.20473137 |
| Thelypodieae | annual | 0.03099634 | 0.00627389 | 0.00654888 | 841.947965 | 840.304406 | 0.09617648 |
| Alysseae | perennial | 0.02644351 | 0.29525668 | 0.01317552 | 996.015901 | 996.561161 | 0.02389206 |
| Antirrhineae | perennial | 0.01566215 | 0.03345322 | 0.00567421 | 837.302126 | 838.907877 | 0.0552575 |
| Apioideae | perennial | 0.01284296 | 0.07987379 | 0.01407058 | 923.416731 | 923.560248 | 0.09002476 |
| Arabideae | perennial | 0.25307988 | 0.68015068 | 0.14579052 | 991.071352 | 991.113327 | 0.10693386 |
| Balsamiaceae | perennial | 0.01369379 | 0.00380646 | 0.00053683 | 294.505224 | 418.545005 | 0.09613556 |
| Brassiceae | perennial | 0.0212001 | 5.02214681 | 0.35268411 | 904.628579 | 903.539404 | 0.03489985 |
| Cardamineae | perennial | 0.03668898 | 0.3267593 | 0.09180805 | 903.092943 | 903.216508 | 0.1406005 |
| Cardueae | perennial | 0.01723469 | 0.09020761 | 0.01433502 | 926.199939 | 926.071184 | 0.06900279 |
| CES | perennial | 0.03064458 | 0.00666191 | 0.00627936 | 790.30862 | 812.204552 | 0.15948445 |
| Chamaecrista | perennial | 0.01191969 | 0.13331923 | 0.00166961 | 417.479165 | 417.536707 | 0.0063186 |
| Croton | perennial | 0.00530538 | 0.13380144 | 0.01444485 | 441.669282 | 449.858644 | 0.05466812 |
| Erysimeae | perennial | 0.0282361 | 0.05325203 | 0.01877336 | 1006.94915 | 1006.95589 | 0.1251606 |
| Euclidieae | perennial | 0.05983 | 0.57957188 | 0.04756543 | 1135.80506 | 1136.15584 | 0.04448762 |
| Eumalvoideae | perennial | 0.01466083 | 0.07993348 | 0.01366449 | 553.671627 | 554.652931 | 0.08879878 |
| Gesneriaceae | perennial | 0.00239728 | 0.00011715 | 0.0027086 | 479.622938 | 479.632455 | 0.19628061 |
| Grewioideae | perennial | 0.00198939 | 0.05615118 | 0.00888641 | 461.762101 | 465.804725 | 0.07974829 |
| Heliophileae | perennial | 0.02053737 | 0.22994972 | 0.00262652 | 676.483309 | 678.326028 | 0.00675669 |
| Hypericum | perennial | 0.0135637 | 0.02634112 | 0.00773588 | 859.501785 | 872.116987 | 0.33904791 |
| Lepidieae | perennial | 0.20604805 | 1.20419534 | 0.17984272 | 809.447138 | 820.311873 | 0.07520574 |
| Lupinus | perennial | 0.08571184 | 0.00150559 | 0.02851749 | 698.858054 | 759.650441 | 0.17487776 |
| Lysimachieae | perennial | 0.02746466 | 0.00206873 | 0.00905691 | 826.014558 | 825.973939 | 0.24500091 |
| Onagraceae | perennial | 0.010763 | 0.05566329 | 0.01796286 | 673.965449 | 682.706142 | 0.23232201 |
| Orobanchaceae | perennial | 0.04493717 | 0.10805047 | 0.02204674 | 878.88469 | 876.584242 | 0.10108514 |
| Panicoideae | perennial | 0.04248433 | 5.82108862 | 0.98241422 | 548.358333 | 548.364392 | 0.08443103 |
| Polemoniaceae | perennial | 0.01019994 | 0.03094406 | 0.00516661 | 418.137332 | 689.649525 | 0.07613777 |
| Pooideae | perennial | 0.0313471 | 0.59969435 | 0.15827396 | 873.220878 | 873.636097 | 0.13148293 |
| Primulaceae | perennial | 0.01169352 | 2.55118575 | 0.19162347 | 940.087717 | 940.121823 | 0.03752795 |
| Rubieae | perennial | 0.01132199 | 0.95176001 | 0.08604369 | 887.015009 | 886.936364 | 0.04423067 |
| Salvia | perennial | 0.01331384 | 0.04632607 | 0.01239111 | 732.177346 | 732.523841 | 0.12671422 |
| Solanaceae | perennial | 0.01325026 | 0.10759233 | 0.0277963 | 527.868206 | 528.177482 | 0.12907516 |
| Spermacoceae | perennial | 0.02360441 | 0.0574132 | 0.01400538 | 494.215488 | 491.398934 | 0.12128527 |
| Thelypodieae | perennial | 0.07839508 | 0.00627389 | 0.00666205 | 839.593379 | 840.144516 | 0.09670056 |

**Table S3** Parameter estimates from the model averaged hOUwie fits for BIO5.

| Group.1 | Group.2 | rates | alpha | sigma.sq | theta | expected_mean | expected_var |
|---|---|---|---|---|---|---|---|
| Alysseae | annual | 0.04337222 | 10.6994061 | 0.00181687 | 27.1166682 | 27.1134443 | 8.84E-05 |
| Antirrhineae | annual | 0.03134976 | 0.10723967 | 3.72E-05 | 29.7492161 | 29.7062696 | 0.00017423 |
| Apioideae | annual | 0.01620089 | 0.19195569 | 9.63E-05 | 31.5150133 | 29.4248292 | 0.00024946 |
| Arabideae | annual | 0.27197746 | 2.01303375 | 0.00133574 | 23.0705727 | 21.9706136 | 0.0003282 |
| Balsamiaceae | annual | 0.01449122 | 0.25096766 | 7.73E-05 | 24.4622633 | 24.4695021 | 0.00015536 |
| Brassiceae | annual | 0.05795414 | 10.2964912 | 0.00437339 | 28.505024 | 28.5050131 | 0.00021431 |
| Cardamineae | annual | 0.02563818 | 6.74858649 | 0.00272862 | 26.0792835 | 25.8167091 | 0.00020681 |
| Cardueae | annual | 0.0714986 | 0.12847278 | 4.84E-05 | 29.6563315 | 29.4250386 | 0.00018871 |
| CES | annual | 0.1047205 | 0.00790863 | 1.86E-05 | 168.146636 | 18.8642562 | 0.00051714 |
| Chamaecrista | annual | 0.09043656 | 0.45939341 | 2.18E-05 | 31.159002 | 30.6792633 | 2.95E-05 |
| Croton | annual | 0.02748687 | 0.11922311 | 2.21E-05 | 29.0554108 | 29.0487245 | 9.27E-05 |
| Erysimeae | annual | 0.18148684 | 4.08360477 | 0.00064567 | 25.4569097 | 25.4175552 | 0.00016716 |
| Euclidieae | annual | 0.03490325 | 13.684164 | 0.02284841 | 28.7276635 | 28.7031489 | 0.00082231 |
| Eumalvoideae | annual | 0.03909404 | 0.06037643 | 2.93E-05 | 28.4445017 | 28.4219481 | 0.00024352 |
| Gesneriaceae | annual | 0.007603 | 0.0490575 | 9.58E-06 | 26.4865403 | 26.4864758 | 9.90E-05 |
| Grewioideae | annual | 0.02291133 | 2.54756316 | 0.00032241 | 30.8505594 | 30.8194556 | 6.42E-05 |
| Heliophileae | annual | 0.0345661 | 3.84160829 | 0.0002021 | 28.2760965 | 28.27605 | 2.42E-05 |
| Hypericum | annual | 0.08139911 | 0.05856605 | 5.05E-05 | 28.2388658 | 26.8724685 | 0.00048208 |
| Lepidieae | annual | 0.27812295 | 12.7990663 | 0.00756354 | 28.9273622 | 28.8702129 | 0.00029957 |
| Lupinus | annual | 0.03112251 | 0.02696097 | 2.58E-05 | 26.4386383 | 26.3680204 | 0.00026135 |
| Lysimachieae | annual | 0.04106105 | 0.12980581 | 2.50E-05 | 31.8765016 | 26.366249 | 0.00010913 |
| Onagraceae | annual | 0.02918668 | 0.47536547 | 0.00040129 | 30.9680118 | 30.7841683 | 0.00042586 |
| Orobanchaceae | annual | 0.04944369 | 0.36440144 | 0.00012831 | 26.9652409 | 26.867926 | 0.00018263 |
| Panicoideae | annual | 0.02191758 | 10.1067817 | 0.0017054 | 30.5397738 | 30.5380222 | 8.52E-05 |
| Polemoniaceae | annual | 0.01169879 | 0.07125677 | 4.05E-05 | 28.8676101 | 28.8663668 | 0.000284 |
| Pooideae | annual | 0.02549196 | 9.27369114 | 0.00549692 | 28.2741001 | 28.2095742 | 0.00029637 |
| Primulaceae | annual | 0.03001064 | 0.22886667 | 0.00011637 | 20.4227465 | 20.2923207 | 0.00025699 |
| Rubieae | annual | 0.07146282 | 11.3357708 | 0.00180116 | 27.3842536 | 27.3710283 | 8.23E-05 |
| Salvia | annual | 0.01791912 | 0.12517184 | 2.55E-05 | 27.4405428 | 27.2627386 | 0.00013596 |
| Solanaceae | annual | 0.03815547 | 0.16530149 | 7.87E-05 | 25.6218183 | 25.6218162 | 0.00023809 |
| Spermacoceae | annual | 0.03353691 | 0.11487522 | 1.19E-05 | 29.1766623 | 29.1766282 | 5.15E-05 |
| Thelypodieae | annual | 0.03237878 | 0.0634584 | 7.00E-05 | 31.7862202 | 29.6540432 | 0.00056777 |
| Alysseae | perennial | 0.02426779 | 10.6994061 | 0.00220883 | 26.6235869 | 26.6237214 | 0.00010143 |
| Antirrhineae | perennial | 0.01203507 | 0.10723967 | 3.72E-05 | 27.4527429 | 28.3186798 | 0.00017427 |
| Apioideae | perennial | 0.01339412 | 0.19195569 | 9.24E-05 | 24.4896495 | 24.5452359 | 0.0002408 |
| Arabideae | perennial | 0.07331012 | 2.01303375 | 0.00130584 | 18.8053626 | 18.8193568 | 0.00032613 |
| Balsamiaceae | perennial | 0.01216597 | 0.25096766 | 7.21E-05 | 24.6623734 | 24.6397144 | 0.00014281 |
| Brassiceae | perennial | 0.0193124 | 10.2964912 | 0.0048282 | 28.4783565 | 28.4786712 | 0.00022954 |
| Cardamineae | perennial | 0.03506271 | 6.74858649 | 0.00289792 | 21.2773619 | 21.2810615 | 0.00021244 |
| Cardueae | perennial | 0.01506848 | 0.12847278 | 4.84E-05 | 24.1898272 | 26.2974977 | 0.0001887 |
| CES | perennial | 0.02988037 | 0.00790863 | 2.07E-05 | 18.4419802 | 18.4764177 | 0.00052526 |
| Chamaecrista | perennial | 0.01294343 | 0.45939341 | 2.57E-05 | 29.0915432 | 29.1208672 | 2.97E-05 |
| Croton | perennial | 0.00572356 | 0.11922311 | 2.21E-05 | 29.0406205 | 29.0407615 | 9.28E-05 |
| Erysimeae | perennial | 0.02578306 | 4.08360477 | 0.00165405 | 25.2774197 | 25.278247 | 0.00020125 |
| Euclidieae | perennial | 0.0509384 | 13.684164 | 0.00071978 | 13.8083493 | 13.8495702 | 5.00E-05 |
| Eumalvoideae | perennial | 0.01214217 | 0.06037643 | 2.90E-05 | 28.4211974 | 28.4212627 | 0.0002419 |
| Gesneriaceae | perennial | 0.00236961 | 0.0490575 | 9.61E-06 | 26.486387 | 26.4863866 | 0.00010048 |
| Grewioideae | perennial | 0.00347057 | 2.54756316 | 0.00020284 | 29.3816285 | 29.3899184 | 4.59E-05 |
| Heliophileae | perennial | 0.02074398 | 3.84160829 | 0.0001305 | 26.9801262 | 27.0460602 | 2.14E-05 |
| Hypericum | perennial | 0.01772965 | 0.05856605 | 4.94E-05 | 26.4522746 | 26.4913315 | 0.00047563 |
| Lepidieae | perennial | 0.23259435 | 12.7990663 | 0.00853369 | 27.1273164 | 27.1996417 | 0.00033105 |
| Lupinus | perennial | 0.08186885 | 0.02696097 | 8.23E-05 | 25.6003891 | 25.8845822 | 0.00050458 |
| Lysimachieae | perennial | 0.02547939 | 0.12980581 | 3.45E-05 | 26.3543955 | 26.3548318 | 0.00013576 |
| Onagraceae | perennial | 0.01116725 | 0.47536547 | 0.00029177 | 25.5988822 | 25.9113943 | 0.00031558 |
| Orobanchaceae | perennial | 0.04580571 | 0.36440144 | 0.0001489 | 26.1907864 | 26.2268877 | 0.00020935 |
| Panicoideae | perennial | 0.0407256 | 10.1067817 | 0.00165495 | 29.6247791 | 29.6269721 | 8.18E-05 |
| Polemoniaceae | perennial | 0.01145944 | 0.07125677 | 4.05E-05 | 28.8512318 | 28.8583737 | 0.00028402 |
| Pooideae | perennial | 0.03070636 | 9.27369114 | 0.00549692 | 21.7025115 | 21.7098626 | 0.00029637 |
| Primulaceae | perennial | 0.01104393 | 0.22886667 | 0.00011736 | 20.0237445 | 20.0274335 | 0.00025704 |
| Rubieae | perennial | 0.02879279 | 11.3357708 | 0.00233061 | 24.2344206 | 24.2428943 | 0.00010212 |
| Salvia | perennial | 0.01330184 | 0.12517184 | 4.68E-05 | 27.1876332 | 27.1876988 | 0.00018666 |
| Solanaceae | perennial | 0.01382659 | 0.16530149 | 7.87E-05 | 25.6218135 | 25.6218136 | 0.00023809 |
| Spermacoceae | perennial | 0.06163745 | 0.11487522 | 1.30E-05 | 29.1761336 | 29.1763155 | 0.0001138 |
| Thelypodieae | perennial | 0.07627419 | 0.0634584 | 7.21E-05 | 27.2564202 | 27.4590681 | 0.00057905 |

**Table S4** Parameter estimates from the model averaged hOUwie fits for BIO6

| Group.1 | Group.2 | rates | alpha | sigma.sq | theta | expected_mean | expected_var |
|---|---|---|---|---|---|---|---|
| Alysseae | annual | 0.05161333 | 6.22818656 | 0.00420731 | -2.5872065 | -2.6130718 | 0.00035728 |
| Antirrhineae | annual | 0.03153751 | 0.05956301 | 4.77E-05 | 4.16352647 | 4.16319215 | 0.00040351 |
| Apioideae | annual | 0.01369082 | 0.11671906 | 5.17E-05 | 2.79702488 | 0.0871027 | 0.00036986 |
| Arabideae | annual | 1.57546449 | 1.02115052 | 0.00222512 | -8.9057323 | -9.3703577 | 0.0010899 |
| Balsamiaceae | annual | 0.01404574 | 0.00331545 | 2.63E-05 | 4.33322988 | 5.0580273 | 0.00160415 |
| Brassiceae | annual | 0.05940327 | 8.37844032 | 0.00515641 | 2.79205995 | 2.79147376 | 0.00031426 |
| Cardamineae | annual | 0.03710529 | 14.9016984 | 0.00896339 | -0.8557734 | -0.9579275 | 0.00031157 |
| Cardueae | annual | 0.07951689 | 0.04745138 | 5.89E-05 | 24.2413272 | 5.22621163 | 0.00075799 |
| CES | annual | 0.08983146 | 0.00061035 | 4.97E-05 | -6.9837056 | -8.0944107 | 0.00108717 |
| Chamaecrista | annual | 0.0776548 | 0.06377404 | 1.31E-05 | 17.7269466 | 17.5173651 | 0.00010645 |
| Croton | annual | 0.04789096 | 0.08200858 | 0.00012471 | 0.35893374 | 5.81302847 | 0.00067711 |
| Erysimeae | annual | 0.18334152 | 0.130955 | 0.00021388 | -5.4326107 | -5.4313851 | 0.00099315 |
| Euclidieae | annual | 0.04332979 | 0.78484052 | 0.00137342 | -9.4705188 | -9.5720676 | 0.00107547 |
| Eumalvoideae | annual | 0.04499692 | 0.0400512 | 4.37E-05 | 10.7729575 | 9.68704456 | 0.00054352 |
| Gesneriaceae | annual | 0.01200571 | 1.80E-05 | 1.86E-05 | 32.2990803 | 11.351057 | 0.00128112 |
| Grewioideae | annual | 0.0188415 | 0.03116208 | 2.40E-05 | 14.1142749 | 14.2676253 | 0.00038977 |
| Heliophileae | annual | 0.03543767 | 0.72214292 | 6.28E-05 | 4.42898155 | 4.42874101 | 6.42E-05 |
| Hypericum | annual | 0.08505912 | 0.38813508 | 0.00039905 | 3.66568345 | 3.60648487 | 0.0005244 |
| Lepidieae | annual | 0.24167275 | 0.15644081 | 0.00047588 | 1.37135046 | 1.53084075 | 0.00201487 |
| Lupinus | annual | 0.0298838 | 8.05E-05 | 1.85E-05 | 2.92253971 | 2.92250733 | 0.00037004 |
| Lysimachieae | annual | 0.03317461 | 0.01845862 | 1.65E-05 | 4.78979231 | -0.226402 | 0.00108976 |
| Onagraceae | annual | 0.0729175 | 0.32771862 | 0.00056161 | 1.74824217 | 1.74069861 | 0.00086319 |
| Orobanchaceae | annual | 0.05366978 | 0.3059429 | 0.00059156 | 0.34673511 | -0.2261732 | 0.00096534 |
| Panicoideae | annual | 0.02806071 | 0.89663704 | 0.00069173 | 11.7845865 | 11.7886472 | 0.00037604 |
| Polemoniaceae | annual | 0.01359697 | 0.03724123 | 3.41E-05 | 0.06272615 | 0.07136543 | 0.00062239 |
| Pooideae | annual | 0.03461644 | 7.60060832 | 0.00550049 | 0.55533901 | 0.46764762 | 0.00036478 |
| Primulaceae | annual | 0.02774153 | 1.80320897 | 0.00309142 | -8.8019942 | -8.8019938 | 0.00075187 |
| Rubieae | annual | 0.12070004 | 14.8097802 | 0.00561357 | 2.15523062 | 2.15510356 | 0.00019313 |
| Salvia | annual | 0.01857804 | 0.08850728 | 7.58E-05 | 4.04848497 | 3.96820796 | 0.00042913 |
| Solanaceae | annual | 0.03417268 | 0.11593327 | 0.00015636 | 7.97218075 | 8.47110923 | 0.00067372 |
| Spermacoceae | annual | 0.03577035 | 0.1393921 | 0.00014899 | 14.4334213 | 14.0807734 | 0.00052781 |
| Thelypodieae | annual | 0.03118705 | 0.76067528 | 0.00024656 | 2.44353163 | 1.62625784 | 0.00014893 |
| Alysseae | perennial | 0.02680039 | 6.22818656 | 0.00449874 | -3.0347174 | -3.0141286 | 0.00036718 |
| Antirrhineae | perennial | 0.01118818 | 0.05956301 | 5.19E-05 | 4.15231742 | 4.15845512 | 0.00042346 |
| Apioideae | perennial | 0.01391388 | 0.11671906 | 0.00019408 | -3.4906913 | -3.4356057 | 0.00082635 |
| Arabideae | perennial | 0.2551859 | 1.02115052 | 0.0022278 | -10.24818 | -10.241348 | 0.00109115 |
| Balsamiaceae | perennial | 0.01467325 | 0.00331545 | 7.37E-05 | 1534.9123 | 9.94250783 | 0.00229979 |
| Brassiceae | perennial | 0.02528941 | 8.37844032 | 0.005576 | 0.97804292 | 0.99501929 | 0.00032824 |
| Cardamineae | perennial | 0.03890158 | 14.9016984 | 0.04083057 | -6.4841025 | -6.4837457 | 0.00136872 |
| Cardueae | perennial | 0.03683252 | 0.04745138 | 0.00011318 | -2.846125 | -0.395766 | 0.00104146 |
| CES | perennial | 0.02897703 | 0.00061035 | 3.21E-05 | -8.0927916 | -8.0954344 | 0.00099018 |
| Chamaecrista | perennial | 0.00983694 | 0.06377404 | 1.32E-05 | 17.395518 | 17.3991579 | 0.0001079 |
| Croton | perennial | 0.01063973 | 0.08200858 | 4.48E-05 | 15.6573085 | 15.274565 | 0.00028631 |
| Erysimeae | perennial | 0.02610273 | 0.130955 | 0.00023434 | -5.4284971 | -5.4285068 | 0.00097366 |
| Euclidieae | perennial | 0.04535073 | 0.78484052 | 0.00170168 | -16.337683 | -15.828586 | 0.00113395 |
| Eumalvoideae | perennial | 0.01453831 | 0.0400512 | 4.36E-05 | 9.67899705 | 9.67982439 | 0.0005434 |
| Gesneriaceae | perennial | 0.00320575 | 1.80E-05 | 1.73E-05 | 18.2306078 | 11.3302853 | 0.00124577 |
| Grewioideae | perennial | 0.00187511 | 0.03116208 | 2.82E-05 | 14.723028 | 14.681001 | 0.00044835 |
| Heliophileae | perennial | 0.02302884 | 0.72214292 | 9.40E-05 | 4.03422465 | 4.17033371 | 6.92E-05 |
| Hypericum | perennial | 0.01869075 | 0.38813508 | 0.00037287 | 3.41154958 | 3.42875815 | 0.00052235 |
| Lepidieae | perennial | 0.20581917 | 0.15644081 | 0.00033798 | 2.06311666 | 1.82117735 | 0.00162877 |
| Lupinus | perennial | 0.07971378 | 8.05E-05 | 0.00020486 | 2.92213576 | 2.92224044 | 0.00116014 |
| Lysimachieae | perennial | 0.02647578 | 0.01845862 | 7.57E-05 | -0.2681844 | -0.2653882 | 0.00142268 |
| Onagraceae | perennial | 0.00338222 | 0.32771862 | 0.00056695 | 2.55685637 | 2.55235806 | 0.0008643 |
| Orobanchaceae | perennial | 0.04425846 | 0.3059429 | 0.00059516 | -3.2859605 | -3.0132383 | 0.00097497 |
| Panicoideae | perennial | 0.04004537 | 0.89663704 | 0.00081318 | 11.8072231 | 11.8068683 | 0.00047584 |
| Polemoniaceae | perennial | 0.01081812 | 0.03724123 | 0.00010675 | 0.22643204 | 0.15817461 | 0.00127746 |
| Pooideae | perennial | 0.03306141 | 7.60060832 | 0.0186809 | -6.3391051 | -6.3288309 | 0.00122822 |
| Primulaceae | perennial | 0.01066659 | 1.80320897 | 0.00261848 | -8.7184746 | -8.7184746 | 0.00072935 |
| Rubieae | perennial | 0.01063675 | 14.8097802 | 0.01562851 | -2.2668226 | -2.2582081 | 0.00052696 |
| Salvia | perennial | 0.01315338 | 0.08850728 | 7.72E-05 | 3.92992893 | 3.93018933 | 0.00043193 |
| Solanaceae | perennial | 0.01372745 | 0.11593327 | 0.00015632 | 9.1404873 | 9.12695341 | 0.0006736 |
| Spermacoceae | perennial | 0.01355562 | 0.1393921 | 0.00013589 | 8.69078007 | 9.93863497 | 0.0004926 |
| Thelypodieae | perennial | 0.08701517 | 0.76067528 | 0.00021697 | -3.8061756 | -3.7637158 | 0.00014794 |

**Table S5** Parameter estimates from the model averaged hOUwie fits for BIO12

| Group.1 | Group.2 | rates | alpha | sigma.sq | theta | expected_mean | expected_var |
|---|---|---|---|---|---|---|---|
| Alysseae | annual | 0.04737243 | 0.20819149 | 0.04821967 | 515.233724 | 520.845008 | 0.11931237 |
| Antirrhineae | annual | 0.03270945 | 0.07240906 | 0.05400553 | 410.490149 | 412.728464 | 0.37246154 |
| Apioideae | annual | 0.01655018 | 0.10902763 | 0.07480959 | 600.605142 | 615.642547 | 0.34309049 |
| Arabideae | annual | 10.5059625 | 10.0883102 | 5.37488864 | 607.404663 | 608.118428 | 0.30136072 |
| Balsamiaceae | annual | 0.01408315 | 0.06439439 | 0.02217034 | 970.496073 | 1060.84788 | 0.17567393 |
| Brassiceae | annual | 0.0589668 | 1.76016765 | 1.31409528 | 492.717113 | 492.584541 | 0.37679678 |
| Cardamineae | annual | 0.03248418 | 0.53738209 | 0.21357371 | 862.892542 | 860.464327 | 0.19984715 |
| Cardueae | annual | 0.07146612 | 0.08485246 | 0.02860879 | 556.030133 | 555.226047 | 0.17498418 |
| CES | annual | 0.10051921 | 0.0897381 | 0.08183852 | 318.204754 | 316.975313 | 0.67019208 |
| Chamaecrista | annual | 0.09882294 | 0.04823854 | 0.00645252 | 1288.22694 | 1285.88771 | 0.10203044 |
| Croton | annual | 0.0657443 | 0.07057981 | 0.19837714 | 1259.94596 | 1263.75014 | 0.96853921 |
| Erysimeae | annual | 0.18059134 | 0.33982324 | 0.1062191 | 441.109953 | 459.592067 | 0.15821803 |
| Euclidieae | annual | 0.04122487 | 0.10512365 | 0.02499272 | 226.192034 | 226.189226 | 0.1351553 |
| Eumalvoideae | annual | 0.05066802 | 0.05628234 | 0.18044721 | 634.539604 | 652.647046 | 1.30019166 |
| Gesneriaceae | annual | 0.00933779 | 0.00586886 | 0.00540275 | 2401.60746 | 1754.24081 | 0.31425156 |
| Grewioideae | annual | 0.02251709 | 0.01408937 | 0.01068491 | 722.609891 | 989.06991 | 0.33630006 |
| Heliophileae | annual | 0.03311223 | 0.19296797 | 0.08350208 | 243.426276 | 243.520266 | 0.24173503 |
| Hypericum | annual | 0.08401337 | 3.0114711 | 0.6776461 | 1216.79566 | 1216.86001 | 0.11949485 |
| Lepidieae | annual | 0.2466082 | 14.5832526 | 6.02475324 | 393.737997 | 393.860982 | 0.20732368 |
| Lupinus | annual | 0.0297662 | 0.06025797 | 0.05970691 | 562.62679 | 625.787037 | 0.40368399 |
| Lysimachieae | annual | 0.02842698 | 0.11347005 | 0.0110182 | 969.797239 | 966.581562 | 0.09253496 |
| Onagraceae | annual | 0.39683951 | 0.02167004 | 0.09018649 | 954.557296 | 952.29553 | 1.42189999 |
| Orobanchaceae | annual | 0.04945723 | 0.07389371 | 0.04647261 | 730.129648 | 706.770783 | 0.33064432 |
| Panicoideae | annual | 0.02474128 | 0.48334241 | 0.247008 | 1035.40849 | 1049.03123 | 0.25616232 |
| Polemoniaceae | annual | 0.01148871 | 0.01947166 | 0.02355574 | 39.6677214 | 143.598729 | 0.53311352 |
| Pooideae | annual | 0.02238177 | 8.59686616 | 2.64449224 | 583.224332 | 583.224332 | 0.15427025 |
| Primulaceae | annual | 0.03370578 | 0.20251442 | 0.00363834 | 424.745044 | 473.130039 | 0.04871923 |
| Rubieae | annual | 0.04086928 | 0.5448742 | 0.01326983 | 577.69731 | 611.146904 | 0.03277425 |
| Salvia | annual | 0.02865751 | 0.05890604 | 1.66478395 | 532.772536 | 567.133825 | 5.73621352 |
| Solanaceae | annual | 0.03071688 | 0.11665591 | 0.19142686 | 511.278857 | 665.148487 | 0.81962612 |
| Spermacoceae | annual | 0.03606006 | 0.05363067 | 0.02618234 | 1238.42687 | 1218.88448 | 0.24925646 |
| Thelypodieae | annual | 0.03025708 | 0.28845223 | 0.29293827 | 656.824222 | 541.813074 | 0.52192279 |
| Alysseae | perennial | 0.02317844 | 0.20819149 | 0.03804949 | 579.780784 | 570.514228 | 0.0983292 |
| Antirrhineae | perennial | 0.01227838 | 0.07240906 | 0.0632659 | 505.82864 | 458.96603 | 0.41929557 |
| Apioideae | perennial | 0.01301137 | 0.10902763 | 0.07482765 | 634.694502 | 634.445528 | 0.34315586 |
| Arabideae | perennial | 1.48285992 | 10.0883102 | 5.4521792 | 613.045861 | 613.041922 | 0.27675403 |
| Balsamiaceae | perennial | 0.01451376 | 0.06439439 | 0.02015306 | 1871.70662 | 1679.55734 | 0.15954815 |
| Brassiceae | perennial | 0.02160555 | 1.76016765 | 1.29222597 | 416.796016 | 420.584472 | 0.36448539 |
| Cardamineae | perennial | 0.03955619 | 0.53738209 | 0.28257672 | 852.276083 | 852.355688 | 0.26755142 |
| Cardueae | perennial | 0.01769078 | 0.08485246 | 0.05928923 | 556.051389 | 555.551978 | 0.2925052 |
| CES | perennial | 0.03118811 | 0.0897381 | 0.10409495 | 316.056617 | 316.120842 | 0.7323502 |
| Chamaecrista | perennial | 0.01088525 | 0.04823854 | 0.00972918 | 1284.65185 | 1284.67505 | 0.11594419 |
| Croton | perennial | 0.00382829 | 0.07057981 | 0.03191024 | 1267.62649 | 1267.47535 | 0.30310253 |
| Erysimeae | perennial | 0.02581131 | 0.33982324 | 0.10191942 | 490.226032 | 489.919729 | 0.15299116 |
| Euclidieae | perennial | 0.04698327 | 0.10512365 | 0.18484741 | 220.197295 | 226.330659 | 0.71344613 |
| Eumalvoideae | perennial | 0.01704625 | 0.05628234 | 0.05717853 | 673.317734 | 670.450098 | 0.58206823 |
| Gesneriaceae | perennial | 0.00225536 | 0.00586886 | 0.00597045 | 1704.92509 | 1693.48979 | 0.32379055 |
| Grewioideae | perennial | 0.0033909 | 0.01408937 | 0.01036084 | 1366.88548 | 1323.32213 | 0.32981363 |
| Heliophileae | perennial | 0.02190553 | 0.19296797 | 0.06775391 | 467.928166 | 385.223352 | 0.21372747 |
| Hypericum | perennial | 0.01755665 | 3.0114711 | 0.23064585 | 1380.56798 | 1380.5384 | 0.03965665 |
| Lepidieae | perennial | 0.22744216 | 14.5832526 | 6.02021216 | 398.185235 | 398.029268 | 0.20592134 |
| Lupinus | perennial | 0.08012825 | 0.06025797 | 0.09324341 | 4292.69077 | 1012.19252 | 0.52254639 |
| Lysimachieae | perennial | 0.02461495 | 0.11347005 | 0.03375462 | 963.492792 | 963.560334 | 0.16066922 |
| Onagraceae | perennial | 0.19986618 | 0.02167004 | 0.13556119 | 956.241676 | 955.842784 | 1.52437791 |
| Orobanchaceae | perennial | 0.04446025 | 0.07389371 | 0.0555584 | 606.228458 | 642.711882 | 0.38035432 |
| Panicoideae | perennial | 0.04163422 | 0.48334241 | 0.24577139 | 1093.43089 | 1092.4182 | 0.25484125 |
| Polemoniaceae | perennial | 0.01023562 | 0.01947166 | 0.0247927 | 1015.12253 | 356.753845 | 0.54782084 |
| Pooideae | perennial | 0.02824956 | 8.59686616 | 5.69766657 | 583.224333 | 583.224333 | 0.33126481 |
| Primulaceae | perennial | 0.00759099 | 0.20251442 | 0.11394444 | 740.85253 | 727.995877 | 0.27999627 |
| Rubieae | perennial | 0.04769792 | 0.5448742 | 0.17420824 | 746.603781 | 743.041502 | 0.15812164 |
| Salvia | perennial | 0.01275457 | 0.05890604 | 0.05316311 | 575.924195 | 575.922976 | 0.83698316 |
| Solanaceae | perennial | 0.01689401 | 0.11665591 | 0.19105004 | 729.618851 | 752.517491 | 0.81823637 |
| Spermacoceae | perennial | 0.01306143 | 0.05363067 | 0.04043872 | 1000.56226 | 1097.78395 | 0.3387305 |
| Thelypodieae | perennial | 0.08492402 | 0.28845223 | 0.27379223 | 339.642254 | 343.354377 | 0.48334722 |

**Table S6** Parameter estimates from the model averaged hOUwie fits for BIO14

| Group.1 | Group.2 | rates | alpha | sigma.sq | theta | expected_mean | expected_var |
|---|---|---|---|---|---|---|---|
| Alysseae | annual | 0.04579259 | 0.47798098 | 0.48552629 | 9.9509342 | 10.0914011 | 0.51430865 |
| Antirrhineae | annual | 0.03415206 | 0.24270092 | 0.51299472 | 6.59550892 | 6.60005843 | 1.05972644 |
| Apioideae | annual | 0.01662286 | 0.13450909 | 0.33430538 | 10.7888516 | 10.5621439 | 1.242724 |
| Arabideae | annual | 1.75095961 | 7.13680715 | 12.3092249 | 13.303918 | 13.4428904 | 0.86253332 |
| Balsamiaceae | annual | 0.01397567 | 0.10438049 | 0.18847685 | 13.2250231 | 14.024651 | 0.91156299 |
| Brassiceae | annual | 0.05477588 | 0.33993062 | 0.6134576 | 8.48687684 | 8.47316861 | 0.9020602 |
| Cardamineae | annual | 0.032016 | 5.10328054 | 7.85332442 | 27.2031202 | 27.1629205 | 0.77100091 |
| Cardueae | annual | 0.06311925 | 0.11819022 | 0.25717259 | 7.93199866 | 8.18382079 | 1.08767445 |
| CES | annual | 0.26964558 | 0.15524033 | 0.18166066 | 9.40973286 | 8.97862281 | 0.83748149 |
| Chamaecrista | annual | 0.08640687 | 0.06312635 | 0.09218532 | 15.4731671 | 15.5129028 | 0.80729012 |
| Croton | annual | 0.14998827 | 0.14082145 | 0.28026217 | 22.3380453 | 21.3961722 | 0.99944581 |
| Erysimeae | annual | 0.17207479 | 0.73359004 | 1.34351187 | 9.79397017 | 9.73528345 | 0.91958202 |
| Euclidieae | annual | 0.04217904 | 0.09874811 | 0.11277588 | 3.36190769 | 3.3638688 | 0.55111196 |
| Eumalvoideae | annual | 0.04172981 | 0.0772997 | 0.15664278 | 4.5435306 | 6.11844567 | 1.01302429 |
| Gesneriaceae | annual | 0.00439606 | 0.02525216 | 0.05538558 | 30.4554998 | 31.247659 | 1.07526829 |
| Grewioideae | annual | 0.017218 | 0.05160354 | 0.11907702 | 5.83801168 | 7.07659271 | 1.15756398 |
| Heliophileae | annual | 0.03554063 | 0.03527601 | 0.045307 | 7.56826034 | 7.56826037 | 0.38233642 |
| Hypericum | annual | 0.09900478 | 0.63522607 | 0.71258968 | 27.9872035 | 28.2061366 | 0.57292021 |
| Lepidieae | annual | 0.27649931 | 1.71090885 | 3.07276909 | 8.29814105 | 8.31073899 | 0.90678868 |
| Lupinus | annual | 0.02734713 | 0.00204536 | 0.12015314 | 8.78293089 | 8.86029821 | 1.65303628 |
| Lysimachieae | annual | 0.04871493 | 0.37000828 | 0.35028624 | 12.2859542 | 13.9443554 | 0.47584819 |
| Onagraceae | annual | 0.06849037 | 2.62375144 | 5.16135105 | 6.73775487 | 6.8316547 | 0.98358232 |
| Orobanchaceae | annual | 0.04351292 | 5.90084035 | 12.11917 | 13.3759506 | 13.3745486 | 1.0299938 |
| Panicoideae | annual | 0.02188394 | 9.91761498 | 15 | 12.7132505 | 12.714649 | 0.75623093 |
| Polemoniaceae | annual | 0.01190267 | 0.08065316 | 0.10943205 | 4.04979968 | 4.29508993 | 0.68488172 |
| Pooideae | annual | 0.02183113 | 7.97966766 | 15 | 9.5025711 | 9.57202901 | 0.93988877 |
| Primulaceae | annual | 0.03226997 | 0.18388311 | 0.45658066 | 9.69604876 | 9.71603307 | 1.24200741 |
| Rubieae | annual | 0.05232785 | 1.17350003 | 2.31212842 | 7.79785992 | 8.49551689 | 0.97091258 |
| Salvia | annual | 0.04507623 | 0.04112045 | 0.12576575 | 8.17490282 | 8.11720532 | 1.42247451 |
| Solanaceae | annual | 0.01689352 | 0.17648081 | 0.42858963 | 6.80269218 | 9.15145633 | 1.18876228 |
| Spermacoceae | annual | 0.03242268 | 0.01405353 | 0.07047834 | 7.15587134 | 10.2375225 | 1.84542428 |
| Thelypodieae | annual | 0.03439999 | 0.0160098 | 0.35180244 | 5.30243797 | 5.05586878 | 2.05078883 |
| Alysseae | perennial | 0.02561623 | 0.47798098 | 0.49790275 | 12.4415605 | 12.3116039 | 0.52624369 |
| Antirrhineae | perennial | 0.01193103 | 0.24270092 | 0.5156674 | 6.98042699 | 6.88094423 | 1.06624138 |
| Apioideae | perennial | 0.01356522 | 0.13450909 | 0.33430401 | 10.2279576 | 10.2320775 | 1.24271999 |
| Arabideae | perennial | 0.27782878 | 7.13680715 | 12.3001201 | 14.2696304 | 14.2675431 | 0.86222939 |
| Balsamiaceae | perennial | 0.01531837 | 0.10438049 | 0.18619607 | 27.860556 | 25.2679439 | 0.89776156 |
| Brassiceae | perennial | 0.02769703 | 0.33993062 | 0.61357049 | 6.66054602 | 6.97911116 | 0.9024216 |
| Cardamineae | perennial | 0.03783889 | 5.10328054 | 7.85333359 | 26.174041 | 26.1743258 | 0.77104292 |
| Cardueae | perennial | 0.03938396 | 0.11819022 | 0.25722806 | 8.61387148 | 8.55311593 | 1.08786262 |
| CES | perennial | 0.04409831 | 0.15524033 | 0.1951037 | 8.51454294 | 8.5419729 | 0.86292059 |
| Chamaecrista | perennial | 0.01085283 | 0.06312635 | 0.09696641 | 15.5104573 | 15.5105721 | 0.86454123 |
| Croton | perennial | 0.03595118 | 0.14082145 | 0.2747635 | 20.5121542 | 20.5238386 | 0.97835386 |
| Erysimeae | perennial | 0.02732867 | 0.73359004 | 1.34366493 | 9.5409786 | 9.54214389 | 0.91991508 |
| Euclidieae | perennial | 0.05597158 | 0.09874811 | 0.11686561 | 3.39611739 | 3.38736917 | 0.56852722 |
| Eumalvoideae | perennial | 0.01216178 | 0.0772997 | 0.15654563 | 9.20691145 | 8.79779358 | 1.01253657 |
| Gesneriaceae | perennial | 0.00249704 | 0.02525216 | 0.05382633 | 31.5242881 | 31.5495104 | 1.0542478 |
| Grewioideae | perennial | 0.00173364 | 0.05160354 | 0.10964436 | 20.5083508 | 19.0210843 | 1.07327493 |
| Heliophileae | perennial | 0.02304538 | 0.03527601 | 0.03994513 | 17.5842476 | 9.9381081 | 0.37277583 |
| Hypericum | perennial | 0.01491198 | 0.63522607 | 0.07844782 | 59.463865 | 57.103958 | 0.06760869 |
| Lepidieae | perennial | 0.21766085 | 1.71090885 | 2.67169604 | 8.43403409 | 8.41705845 | 0.78448221 |
| Lupinus | perennial | 0.06808381 | 0.00204536 | 0.40852831 | 11.4604872 | 9.12873959 | 2.80868967 |
| Lysimachieae | perennial | 0.02524198 | 0.37000828 | 0.35031334 | 21.3256514 | 21.1974096 | 0.47590472 |
| Onagraceae | perennial | 0.00193533 | 2.62375144 | 5.16135105 | 24.7771196 | 24.7621258 | 0.98358232 |
| Orobanchaceae | perennial | 0.04431145 | 5.90084035 | 12.126715 | 13.3227365 | 13.322738 | 1.03114656 |
| Panicoideae | perennial | 0.03429727 | 9.91761498 | 15 | 16.5899866 | 16.5904226 | 0.75623093 |
| Polemoniaceae | perennial | 0.01170706 | 0.08065316 | 0.14545828 | 14.8172803 | 10.4108599 | 0.87436157 |
| Pooideae | perennial | 0.03047146 | 7.97966766 | 15 | 17.0460389 | 17.0333096 | 0.93988877 |
| Primulaceae | perennial | 0.01208718 | 0.18388311 | 0.45738327 | 9.741042 | 9.74052118 | 1.24384541 |
| Rubieae | perennial | 0.04250172 | 1.17350003 | 2.36764626 | 16.1437052 | 15.9928824 | 1.01511516 |
| Salvia | perennial | 0.01256396 | 0.04112045 | 0.12576802 | 8.08740665 | 8.09385121 | 1.42252247 |
| Solanaceae | perennial | 0.01383322 | 0.17648081 | 0.36442223 | 18.2393939 | 18.0283888 | 1.03435575 |
| Spermacoceae | perennial | 0.01776849 | 0.01405353 | 0.07153936 | 11.3450197 | 11.2982585 | 1.86078232 |
| Thelypodieae | perennial | 0.08706407 | 0.0160098 | 0.03774245 | 5.0502574 | 5.05519723 | 0.60973126 |

**Table S7** Parameter estimates from the model averaged hOUwie fits for BIO15

| Group.1 | Group.2 | rates | alpha | sigma.sq | theta | expected_mean | expected_var |
|---|---|---|---|---|---|---|---|
| Alysseae | annual | 0.04378608 | 2.13850345 | 0.46763675 | 44.6338786 | 44.6038733 | 0.11262597 |
| Antirrhineae | annual | 0.03372723 | 0.22175491 | 0.07540967 | 51.1135599 | 51.1235697 | 0.16993306 |
| Apioideae | annual | 0.01412433 | 0.11162976 | 0.06055915 | 45.4198392 | 47.5607071 | 0.27139224 |
| Arabideae | annual | 0.65368158 | 8.94282157 | 3.42358735 | 45.9269541 | 45.9181889 | 0.19142823 |
| Balsamiaceae | annual | 0.01491422 | 0.05322773 | 0.01526969 | 63.9284512 | 63.2497237 | 0.14428678 |
| Brassiceae | annual | 0.05721943 | 1.24043227 | 0.67799626 | 45.5538977 | 45.5556989 | 0.27514224 |
| Cardamineae | annual | 0.02936977 | 0.43801139 | 0.17261584 | 35.9267652 | 36.0347586 | 0.20407219 |
| Cardueae | annual | 0.05806122 | 0.18148917 | 0.04698407 | 48.1409986 | 48.4722471 | 0.12952768 |
| CES | annual | 0.09189669 | 0.01040016 | 0.0134917 | 54.8359681 | 55.0637738 | 0.42128873 |
| Chamaecrista | annual | 0.09217477 | 0.00349425 | 0.00627452 | 57.6550366 | 48.2869124 | 0.1696715 |
| Croton | annual | 0.02436305 | 0.15985228 | 0.06077946 | 32.3514008 | 39.0265804 | 0.1832457 |
| Erysimeae | annual | 0.18234183 | 2.02076521 | 0.82082664 | 45.8866138 | 46.0451014 | 0.18591706 |
| Euclidieae | annual | 0.04168759 | 0.09995358 | 0.06692274 | 63.2240541 | 63.2284362 | 0.47444223 |
| Eumalvoideae | annual | 0.0491412 | 0.09137134 | 0.00616466 | 69.6805507 | 69.1502102 | 0.05506522 |
| Gesneriaceae | annual | 0.00358917 | 0.04584437 | 0.01611679 | 54.6190547 | 54.5101167 | 0.17579113 |
| Grewioideae | annual | 0.01976282 | 0.06982646 | 0.00766736 | 80.3547339 | 77.7308253 | 0.05562106 |
| Heliophileae | annual | 0.03680605 | 0.14277521 | 0.04783703 | 45.2762252 | 45.2757689 | 0.30145577 |
| Hypericum | annual | 0.10033518 | 0.21882122 | 0.12442934 | 27.6950926 | 27.3387598 | 0.27752427 |
| Lepidieae | annual | 0.24903137 | 2.29896517 | 1.06967053 | 47.7279663 | 47.5693626 | 0.23158558 |
| Lupinus | annual | 0.02967714 | 0.04093387 | 0.03023619 | 46.6765906 | 46.6737109 | 0.28734436 |
| Lysimachieae | annual | 0.03009999 | 0.12299691 | 0.00949672 | 61.7952608 | 52.1695755 | 0.09018851 |
| Onagraceae | annual | 0.07922162 | 0.23355981 | 0.14765297 | 63.1929697 | 63.1855445 | 0.31580209 |
| Orobanchaceae | annual | 0.04711804 | 0.37223064 | 0.19990937 | 46.5125465 | 46.7670322 | 0.268549 |
| Panicoideae | annual | 0.02090337 | 9.51001499 | 3.00651428 | 65.7785569 | 65.7257723 | 0.15807046 |
| Polemoniaceae | annual | 0.01127168 | 0.06905906 | 0.01742792 | 53.5377775 | 53.5221929 | 0.1329685 |
| Pooideae | annual | 0.02274741 | 7.86548183 | 3.74703147 | 46.5796933 | 46.4778832 | 0.23815354 |
| Primulaceae | annual | 0.0227563 | 0.09761794 | 0.02988087 | 58.9528172 | 58.703151 | 0.20198982 |
| Rubieae | annual | 0.07596755 | 0.63042086 | 0.22932493 | 49.2887668 | 48.8010052 | 0.17998085 |
| Salvia | annual | 0.02433589 | 0.1049853 | 0.00563725 | 65.2748799 | 64.351494 | 0.09464866 |
| Solanaceae | annual | 0.03319608 | 0.22093706 | 0.09399433 | 61.1618086 | 59.6182792 | 0.21087085 |
| Spermacoceae | annual | 0.03715827 | 0.06165751 | 0.0224194 | 67.3608595 | 66.9352549 | 0.18361977 |
| Thelypodieae | annual | 0.02845831 | 0.02554676 | 0.03513655 | 59.9337805 | 59.1609533 | 0.38825591 |
| Alysseae | perennial | 0.02355089 | 2.13850345 | 0.46343081 | 44.1298154 | 44.1470453 | 0.10804074 |
| Antirrhineae | perennial | 0.01247936 | 0.22175491 | 0.08015895 | 51.8470419 | 51.661791 | 0.18138313 |
| Apioideae | perennial | 0.01350335 | 0.11162976 | 0.06057906 | 50.3990329 | 50.3670744 | 0.2714637 |
| Arabideae | perennial | 0.14104669 | 8.94282157 | 3.42358789 | 45.8690971 | 45.8688795 | 0.19142835 |
| Balsamiaceae | perennial | 0.01234129 | 0.05322773 | 0.01631923 | 57.2302199 | 58.789703 | 0.15279054 |
| Brassiceae | perennial | 0.02125496 | 1.24043227 | 0.54310231 | 46.0897561 | 46.0416265 | 0.22025939 |
| Cardamineae | perennial | 0.03796391 | 0.43801139 | 0.23265672 | 36.2660851 | 36.2622901 | 0.26891698 |
| Cardueae | perennial | 0.0196643 | 0.18148917 | 0.04748735 | 49.9004093 | 49.7022049 | 0.13089321 |
| CES | perennial | 0.02919222 | 0.01040016 | 0.01954551 | 55.1218605 | 55.0930156 | 0.45449213 |
| Chamaecrista | perennial | 0.01116501 | 0.00349425 | 0.00539632 | 48.594263 | 48.57248 | 0.17337735 |
| Croton | perennial | 0.00498416 | 0.15985228 | 0.05019135 | 57.7880607 | 57.4827946 | 0.15721404 |
| Erysimeae | perennial | 0.02781484 | 2.02076521 | 0.85160484 | 48.0497799 | 48.0463584 | 0.21413506 |
| Euclidieae | perennial | 0.04373934 | 0.09995358 | 0.09841652 | 63.3699991 | 63.3476004 | 0.57480906 |
| Eumalvoideae | perennial | 0.01325843 | 0.09137134 | 0.0256599 | 68.352852 | 68.4466995 | 0.13614075 |
| Gesneriaceae | perennial | 0.00254688 | 0.04584437 | 0.01615284 | 54.3060305 | 54.2927852 | 0.176156 |
| Grewioideae | perennial | 0.00180731 | 0.06982646 | 0.01823636 | 55.035188 | 56.4576795 | 0.12787361 |
| Heliophileae | perennial | 0.02387937 | 0.14277521 | 0.07500505 | 45.1016221 | 45.1560854 | 0.34996531 |
| Hypericum | perennial | 0.01532457 | 0.21882122 | 0.04701878 | 25.5684266 | 25.9791231 | 0.12917329 |
| Lepidieae | perennial | 0.21589796 | 2.29896517 | 0.98079118 | 46.280477 | 46.4951824 | 0.21619924 |
| Lupinus | perennial | 0.07455664 | 0.04093387 | 0.11066089 | 44.7449689 | 46.6546953 | 0.56394118 |
| Lysimachieae | perennial | 0.02615461 | 0.12299691 | 0.04818991 | 44.9787417 | 45.138593 | 0.19503272 |
| Onagraceae | perennial | 0.00342103 | 0.23355981 | 0.21365964 | 62.8869461 | 62.9021519 | 0.47123427 |
| Orobanchaceae | perennial | 0.04214628 | 0.37223064 | 0.19945829 | 48.2175298 | 48.1151881 | 0.26801508 |
| Panicoideae | perennial | 0.04139276 | 9.51001499 | 3.00651428 | 54.542389 | 54.5628577 | 0.15807046 |
| Polemoniaceae | perennial | 0.0110541 | 0.06905906 | 0.04797808 | 53.2298641 | 53.3349363 | 0.30786256 |
| Pooideae | perennial | 0.03194115 | 7.86548183 | 3.74729576 | 39.1310502 | 39.1398069 | 0.23832517 |
| Primulaceae | perennial | 0.02759849 | 0.09761794 | 0.07577813 | 58.3878572 | 58.3986934 | 0.27562567 |
| Rubieae | perennial | 0.03582593 | 0.63042086 | 0.26169783 | 46.4002425 | 46.4515327 | 0.21011538 |
| Salvia | perennial | 0.01389107 | 0.1049853 | 0.03814369 | 64.0361411 | 64.0393556 | 0.18099799 |
| Solanaceae | perennial | 0.01338455 | 0.22093706 | 0.08263792 | 54.9069813 | 54.9853456 | 0.18712184 |
| Spermacoceae | perennial | 0.01284506 | 0.06165751 | 0.02612719 | 60.9882376 | 63.540684 | 0.20485035 |
| Thelypodieae | perennial | 0.08355172 | 0.02554676 | 0.03295988 | 58.5349776 | 58.6836054 | 0.37862128 |

**Table S8** Parameter estimates from the model averaged hOUwie fits for AI

| Group.1 | Group.2 | rates | alpha | sigma.sq | theta | expected_mean | expected_var |
|---|---|---|---|---|---|---|---|
| Alysseae | annual | 0.04741912 | 0.38185637 | 0.16395818 | 0.38496298 | 0.38680036 | 0.21213213 |
| Antirrhineae | annual | 0.03300114 | 0.05592467 | 0.06637453 | 0.24546177 | 0.24757722 | 0.59739377 |
| Apioideae | annual | 0.01272887 | 0.08639321 | 0.09000746 | 0.32564161 | 0.4333541 | 0.52083635 |
| Arabideae | annual | 6.71605987 | 13.2761943 | 6.84479096 | 0.56077937 | 0.57307342 | 0.25928284 |
| Balsamiaceae | annual | 0.01514862 | 0.4447766 | 0.11573094 | 0.91051857 | 0.91945291 | 0.12988799 |
| Brassiceae | annual | 0.05482146 | 4.47560605 | 5.86102179 | 0.32844235 | 0.32841307 | 0.65533017 |
| Cardamineae | annual | 0.0263482 | 7.88857953 | 2.49072211 | 0.75645611 | 0.76613676 | 0.15832056 |
| Cardueae | annual | 0.07430414 | 0.12069729 | 0.07107648 | 0.40347295 | 0.40343534 | 0.3026533 |
| CES | annual | 0.1055719 | 0.2152754 | 0.17490966 | 0.23474495 | 0.235634 | 0.45585831 |
| Chamaecrista | annual | 0.09298248 | 0.01037084 | 0.00544259 | 0.84222147 | 0.84571574 | 0.17153597 |
| Croton | annual | 0.3487838 | 0.09015602 | 0.51193033 | 0.68061433 | 0.73109794 | 1.9698587 |
| Erysimeae | annual | 0.1641971 | 0.46829282 | 0.25067817 | 0.37996341 | 0.3808037 | 0.27214973 |
| Euclidieae | annual | 0.04172836 | 0.83170587 | 0.25643256 | 0.15109421 | 0.15160889 | 0.29402523 |
| Eumalvoideae | annual | 0.44474892 | 0.06583692 | 0.14033964 | 0.33734537 | 0.3591446 | 1.03197494 |
| Gesneriaceae | annual | 0.00282602 | 0.03530129 | 0.0039124 | 1.50023931 | 1.29835025 | 0.08969555 |
| Grewioideae | annual | 0.01842587 | 0.02340999 | 0.02914529 | 0.43346657 | 0.56883783 | 0.5637227 |
| Heliophileae | annual | 0.03480187 | 0.29542962 | 0.15327667 | 0.11494827 | 0.11498846 | 0.29686255 |
| Hypericum | annual | 0.08828198 | 0.51092575 | 0.25970734 | 0.9175962 | 0.9246502 | 0.24900597 |
| Lepidieae | annual | 0.22317806 | 13.3117024 | 9.95676983 | 0.23365704 | 0.23396872 | 0.37664908 |
| Lupinus | annual | 0.03089709 | 0.03150713 | 0.08740679 | 0.36252961 | 0.40667594 | 0.76877484 |
| Lysimachieae | annual | 0.0446519 | 0.12977973 | 0.02745905 | 0.66690291 | 0.73395597 | 0.10570229 |
| Onagraceae | annual | 0.09629999 | 0.01888626 | 0.22306228 | 1.58921687 | 0.74529052 | 3.00112032 |
| Orobanchaceae | annual | 0.04708388 | 0.1313756 | 0.14272336 | 0.52534545 | 0.5190375 | 0.55209252 |
| Panicoideae | annual | 0.02900031 | 3.39489086 | 2.51189256 | 0.57060973 | 0.57204269 | 0.36995191 |
| Polemoniaceae | annual | 0.01167439 | 0.03120088 | 0.03792434 | 0.14400596 | 0.14955336 | 0.58897847 |
| Pooideae | annual | 0.0202752 | 2.3078675 | 1.82053275 | 0.38629531 | 0.39451518 | 0.39445122 |
| Primulaceae | annual | 0.03031071 | 0.26184249 | 0.15220615 | 0.62981548 | 0.64549154 | 0.29090594 |
| Rubieae | annual | 0.0437942 | 1.9529575 | 0.74208245 | 0.41998133 | 0.43302712 | 0.18989197 |
| Salvia | annual | 0.02424448 | 0.04983273 | 0.04265632 | 0.22757194 | 0.32798969 | 0.64320078 |
| Solanaceae | annual | 0.0212215 | 0.08909914 | 0.21344954 | 0.47464643 | 0.49185838 | 1.2711778 |
| Spermacoceae | annual | 0.03614699 | 0.03330834 | 0.03165154 | 0.66272183 | 0.65808407 | 0.46702192 |
| Thelypodieae | annual | 0.02820513 | 0.08249061 | 0.16148059 | 0.27164384 | 0.25040851 | 0.95179418 |
| Alysseae | perennial | 0.0259828 | 0.38185637 | 0.19055552 | 0.41248532 | 0.41049499 | 0.25207993 |
| Antirrhineae | perennial | 0.01305015 | 0.05592467 | 0.09786566 | 0.33171154 | 0.27992355 | 0.79199162 |
| Apioideae | perennial | 0.01339697 | 0.08639321 | 0.09000812 | 0.56104836 | 0.55853994 | 0.52083972 |
| Arabideae | perennial | 0.94544963 | 13.2761943 | 6.85569163 | 0.67105727 | 0.67083895 | 0.25941705 |
| Balsamiaceae | perennial | 0.01332795 | 0.4447766 | 0.11650805 | 1.28510119 | 1.26085942 | 0.13349789 |
| Brassiceae | perennial | 0.01996196 | 4.47560605 | 5.85491488 | 0.29661768 | 0.29756153 | 0.65066565 |
| Cardamineae | perennial | 0.03674209 | 7.88857953 | 2.48887785 | 0.92347171 | 0.92330596 | 0.15719276 |
| Cardueae | perennial | 0.01594176 | 0.12069729 | 0.14179715 | 0.4034354 | 0.40343542 | 0.5213694 |
| CES | perennial | 0.03109798 | 0.2152754 | 0.17711983 | 0.23702377 | 0.23697127 | 0.45370278 |
| Chamaecrista | perennial | 0.01308412 | 0.01037084 | 0.00718498 | 0.84640334 | 0.84637807 | 0.18420053 |
| Croton | perennial | 0.00520383 | 0.09015602 | 0.04584149 | 0.7706868 | 0.76631665 | 0.41315199 |
| Erysimeae | perennial | 0.02629642 | 0.46829282 | 0.2540812 | 0.3826006 | 0.38258291 | 0.27579276 |
| Euclidieae | perennial | 0.04985159 | 0.83170587 | 1.69564046 | 0.28977824 | 0.28108604 | 1.0628657 |
| Eumalvoideae | perennial | 0.22596614 | 0.06583692 | 0.11351424 | 0.36789401 | 0.36647704 | 0.88562114 |
| Gesneriaceae | perennial | 0.0024349 | 0.03530129 | 0.01030294 | 1.16120073 | 1.15773544 | 0.14694208 |
| Grewioideae | perennial | 0.00206782 | 0.02340999 | 0.01917063 | 0.84083463 | 0.81014619 | 0.41462109 |
| Heliophileae | perennial | 0.01954472 | 0.29542962 | 0.14374822 | 0.23146351 | 0.19360034 | 0.27538873 |
| Hypericum | perennial | 0.01477116 | 0.51092575 | 0.02165941 | 0.96028264 | 0.95859674 | 0.02420785 |
| Lepidieae | perennial | 0.18862287 | 13.3117024 | 9.94213578 | 0.24430586 | 0.24389481 | 0.37283729 |
| Lupinus | perennial | 0.07555048 | 0.03150713 | 0.12334068 | 38.3543687 | 0.74860672 | 0.90423352 |
| Lysimachieae | perennial | 0.02658251 | 0.12977973 | 0.02650575 | 0.8319011 | 0.8271431 | 0.10307907 |
| Onagraceae | perennial | 0.08027997 | 0.01888626 | 0.10015971 | 0.91257906 | 0.66438018 | 2.27520612 |
| Orobanchaceae | perennial | 0.04453919 | 0.1313756 | 0.14391632 | 0.49402435 | 0.49905313 | 0.56662628 |
| Panicoideae | perennial | 0.04008822 | 3.39489086 | 2.51189256 | 0.65593639 | 0.65522354 | 0.36995191 |
| Polemoniaceae | perennial | 0.01162336 | 0.03120088 | 0.03289652 | 1.29437833 | 0.34390264 | 0.54184284 |
| Pooideae | perennial | 0.02749956 | 2.3078675 | 1.82053275 | 0.59697062 | 0.59608758 | 0.39445122 |
| Primulaceae | perennial | 0.01024173 | 0.26184249 | 0.15969248 | 0.69025438 | 0.6894189 | 0.30677686 |
| Rubieae | perennial | 0.05130808 | 1.9529575 | 0.74247167 | 0.63864692 | 0.63598294 | 0.19018348 |
| Salvia | perennial | 0.01331045 | 0.04983273 | 0.06354232 | 0.35594196 | 0.35594142 | 0.61247128 |
| Solanaceae | perennial | 0.01452606 | 0.08909914 | 0.21376596 | 0.51286719 | 0.51241156 | 1.19297076 |
| Spermacoceae | perennial | 0.01183425 | 0.03330834 | 0.03574066 | 0.59469925 | 0.63138227 | 0.49608885 |
| Thelypodieae | perennial | 0.07986193 | 0.08249061 | 0.12528938 | 0.22268299 | 0.22625942 | 0.81137263 |
